# Cannabidiol Inhibits SARS-CoV-2 Replication and Promotes the Host Innate Immune Response

**DOI:** 10.1101/2021.03.10.432967

**Authors:** Long Chi Nguyen, Dongbo Yang, Vlad Nicolaescu, Thomas J. Best, Takashi Ohtsuki, Shao-Nong Chen, J. Brent Friesen, Nir Drayman, Adil Mohamed, Christopher Dann, Diane Silva, Haley Gula, Krysten A. Jones, J. Michael Millis, Bryan C. Dickinson, Savaş Tay, Scott A. Oakes, Guido F. Pauli, David O. Meltzer, Glenn Randall, Marsha Rich Rosner

## Abstract

The rapid spread of COVID-19 underscores the need for new treatments. Here we report that cannabidiol (CBD), a compound produced by the cannabis plant, inhibits SARS-CoV-2 infection. CBD and its metabolite, 7-OH-CBD, but not congeneric cannabinoids, potently block SARS-CoV-2 replication in lung epithelial cells. CBD acts after cellular infection, inhibiting viral gene expression and reversing many effects of SARS-CoV-2 on host gene transcription. CBD induces interferon expression and up-regulates its antiviral signaling pathway. A cohort of human patients previously taking CBD had significantly lower SARS-CoV-2 infection incidence of up to an order of magnitude relative to matched pairs or the general population. This study highlights CBD, and its active metabolite, 7-OH-CBD, as potential preventative agents and therapeutic treatments for SARS-CoV-2 at early stages of infection.

---

Severe acute respiratory syndrome coronavirus 2 (SARS-CoV-2) is responsible for coronavirus disease 2019 (COVID-19), a pandemic that has overtaken the world during the past year. SARS-CoV-2, related to severe acute respiratory syndrome-related coronavirus (SARS-CoV), is the seventh species of coronavirus known to infect people. These coronaviruses, which include SARS-CoV, 229E, NL63, OC43, HKU1, and MERS-CoV cause a range of symptoms from the common cold to more severe pathologies (*1*). Despite recent vaccine availability, SARS-CoV-2 is still spreading rapidly (*2*), highlighting the need for alternative treatments, especially for populations with limited access to vaccines. To date, few therapies have been identified that block SARS-CoV-2 replication and viral production.

SARS-CoV-2 is a positive-sense single-stranded RNA (+ssRNA) enveloped virus composed of a lipid bilayer and four structural proteins that drive viral particle formation. The spike (S), membrane (M), and envelope (E) are integral proteins of the virus membrane and serve to drive virion budding, while also recruiting the nucleocapsid (N) protein and the viral genomic RNA into nascent virions. Like SARS-CoV, SARS-CoV-2 primarily enters human cells by the binding of the viral S protein to the angiotensin converting enzyme 2 (ACE2) receptor (*3–5*), after which the S protein undergoes proteolysis by transmembrane protease, serine 2 (TMPRSS2) or other proteases into two non-covalently bound peptides (S1, S2) that facilitate viral entry into the host cell. The *N*-terminal S1 binds the ACE2 receptor, and the *C*-terminal S2 mediates viral-cell membrane fusion following proteolytic cleavage by TMRSS2 or other proteases. Depending upon the cell type, viral entry can also occur after ACE2 binding, independent of proteolytic cleavage (*6–8*). Following cell entry, the SARS-CoV-2 genome is translated into two large polypeptides that are cleaved by two viral proteases, MPro and PLPro (*9, 10*), to produce 15 proteins, in addition to the synthesis of subgenomic RNAs that encode another 10 accessory proteins plus the 4 structural proteins. These proteins enable viral replication, assembly, and budding. In an effort to suppress infection by the SARS-CoV-2 beta-coronavirus as well as other evolving pathogenic viruses, we tested the antiviral potential of a number of small molecules that target host stress response pathways.

One potential regulator of the host stress and antiviral inflammatory responses is cannabidiol (CBD), a member of the cannabinoid class of natural products (*11*). CBD is produced by *Cannabis sativa* (Cannabaceae; marijuana/hemp). Hemp refers to cannabis plants or materials derived thereof that contain 0.3% or less of the psychotropic tetrahydrocannabinol (THC) and typically have relatively high CBD content. By contrast, marijuana refers to *C. sativa* materials with more than 0.3% THC by dry weight. THC acts through binding to the cannabinoid receptor, and CBD potentiates this interaction (*12*). Despite numerous studies and many typically unsubstantiated claims related to CBD-containing products, the biology of CBD itself is unclear and specific targets are mostly unknown (*11*). However, an oral solution of CBD is an FDA-approved drug, largely for the treatment of epilepsy (*13*). Thus, CBD has drug status, is viable as a therapeutic, and cannot be marketed as a dietary supplement in the United States (*11*). Although limited, some studies have reported that certain cannabinoids have antiviral effects against hepatitis C virus (HCV) and other viruses (*14*).

## RESULTS

To test the effect of CBD on SARS-CoV-2 replication, we pretreated A549 human lung carcinoma cells expressing exogenous human ACE-2 receptor (A549-ACE2) for 2 hours with 0-10 μM CBD prior to infection with SARS-CoV-2. After 48 hours, we monitored cells for expression of the viral spike protein (S). For comparison, we also treated cells over a similar dose range with an MLK inhibitor (URMC-099) previously implicated as an antiviral for HIV (*12*) and KPT-9274, a PAK4/NAMPT inhibitor (*13*) that our analysis suggested might reverse many changes in gene expression caused by SARS-CoV-2. All three inhibitors potently inhibited viral replication under non-toxic conditions with EC50s ranging from 0.2-2.1 μM (Fig. 1A). CBD inhibited SARS-CoV-2 replication in Vero E6 monkey kidney epithelial cells as well (fig. S1A). No toxicity was observed at the effective doses (fig. S1B). We also determined that CBD suppressed replication of a related beta-coronavirus, mouse hepatitis virus (MHV), under non-toxic conditions with an EC50 of ~5 μM using A549 cells that express the MHV receptor (A549-MHVR), indicating the potential for more broader viral efficacy (fig. S1C,D).

**Fig. 1.**
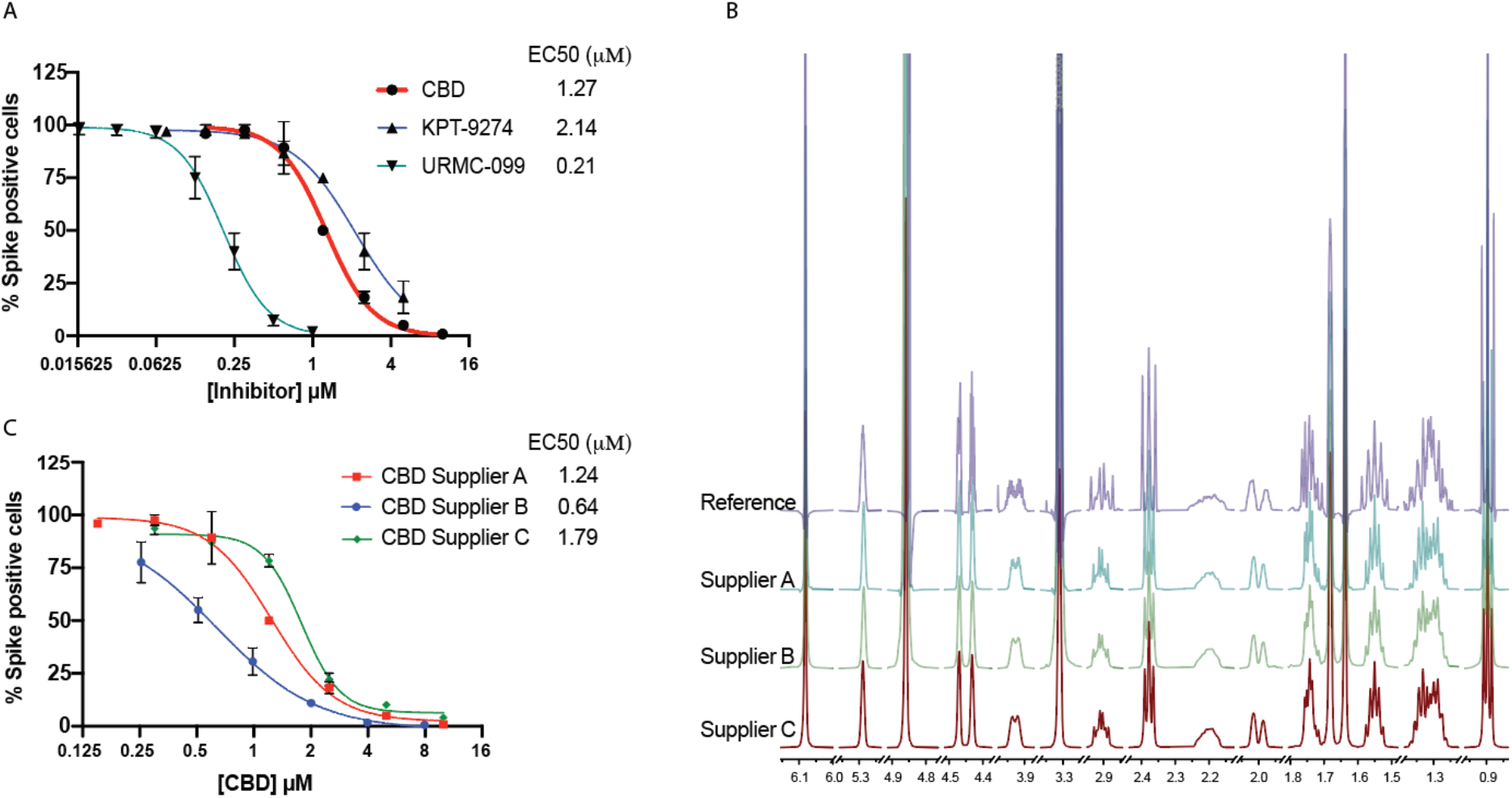
Cannabidiol (CBD) is a potent inhibitor of SARS-CoV-2 infection *in vitro*. (A) A549 cells with ACE2 overexpression (A549-ACE2) were treated with indicated doses of CBD, KPT-9274, or URMC-099 followed by infection with SARS-CoV-2 at a multiplicity of infection (MOI) of 0.5 for 48 hours. The cells were stained for spike protein and the percentage of cells expressing the spike protein in each condition was plotted. EC50 values are indicated. (B) The ^1^H qNMR spectra of CBD from a reference material and CBD samples from three different suppliers A, B, and C. (C) A549-ACE2 cells were treated with indicated doses of CBD from three different suppliers followed by infection with SARS-CoV-2 at an MOI of 0.5 for 48 hours. The cells were stained for spike protein and the percentage of cells expressing the spike protein in each condition was plotted. EC50 values are indicated.

When isolated from its source plant, natural non-synthetic CBD is typically extracted along with other cannabinoids, representing the unavoidable residual complexity of natural products. To verify that CBD is indeed responsible for the viral inhibition, we analyzed a CBD reference standard as well as CBD from three different sources for purity using 100% quantitative NMR (qNMR). These sources included two chemical vendors (Suppliers A and B) and one commercial vendor that used natural materials (Supplier C). The striking congruence between the experimental ^1^H NMR and the recently established quantum-mechanical HiFSA (^1^H Iterative Full Spin Analysis) profiles observed for all materials confirmed that 1) the compounds used were indeed CBD with purities of at least 97% (Fig. 1B) and 2) congeneric cannabinoids were not present at levels above 1.0% (*11*). Analysis of these different CBD preparations in the viral A549-ACE2 infection assay showed similar EC50s with a range from 0.6-1.8 μM likely reflecting the intrinsic biological variability of the assay (Fig. 1C). No toxicity was observed for any of the CBD preparations at the doses used to inhibit viral infection (fig. S1 E-G).

CBD is often consumed as part of a *C. sativa* extract, particularly in combination with psychoactive THC enriched in marijuana plants. We therefore determined whether congeneric cannabinoids, especially analogues with closely related structures and polarities produced by the hemp plant, are also capable of inhibiting SARS-CoV-2 infection. Remarkably, only CBD was a potent agent, while limited or no antiviral activity was exhibited by the structurally closely related congeners that share biosynthesis pathways and form the biogenetically determined residual complexity of CBD purified from *C. sativa*: THC, cannabidiolic acid (CBDA), cannabidivarin (CBDV), cannabichromene (CBC), or cannabigerol (CBG) (Fig. 2 A-B; see Methods). None of these compounds were toxic to the A549-ACE2 cells in the dose range of interest (fig. S2). Notably, combining CBD with THC (1:1) significantly suppressed CBD efficacy consistent with competitive inhibition by THC.

**Fig. 2.**
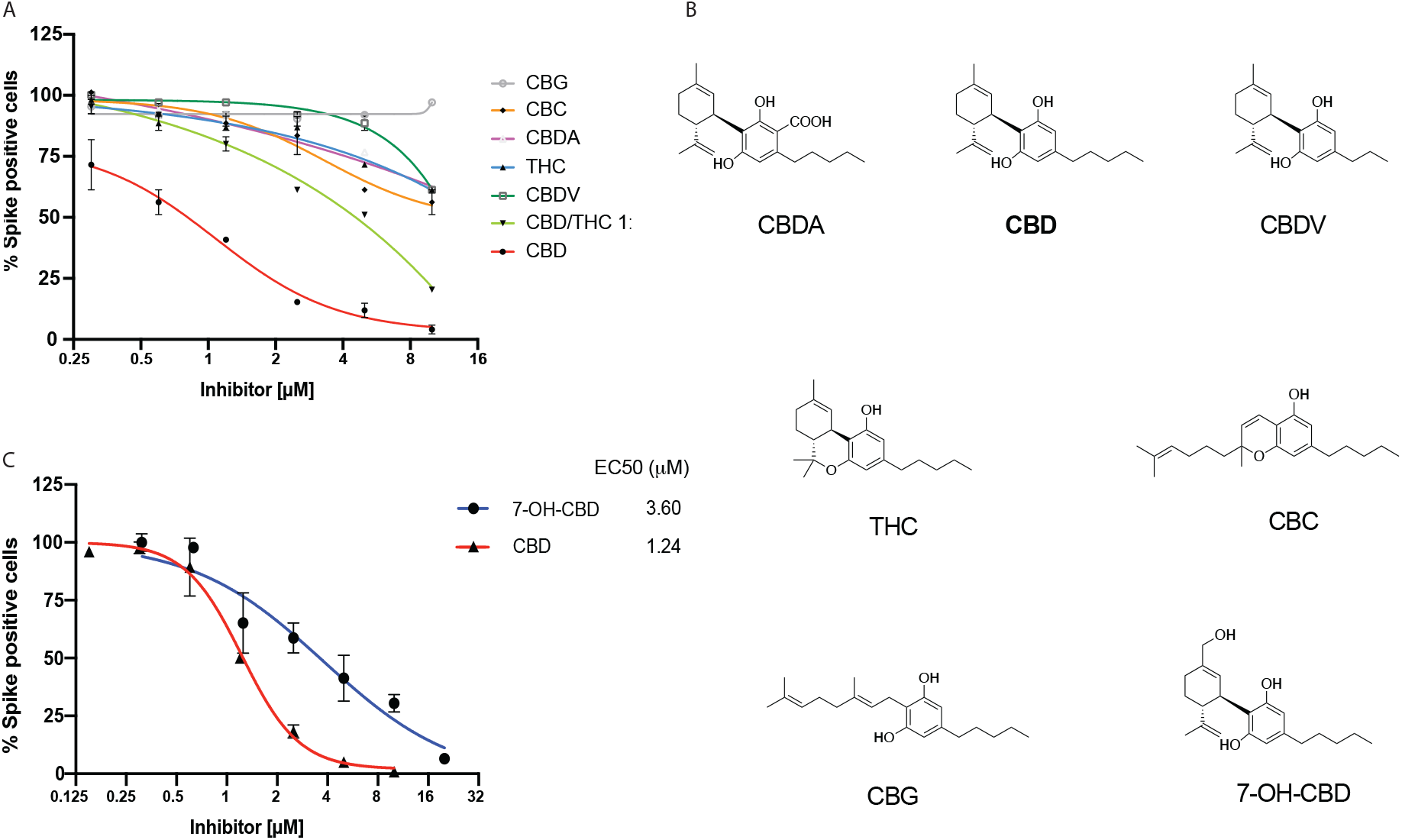
Limited or no inhibition of SARS-CoV-2 infection by Cannabinoids other than CBD. (A) A549-ACE2 cells were treated with indicated doses of various cannabinoids or a CBD/THC 1:1 mixture followed by infection with SARS-CoV-2 at an MOI of 0.5 for 48 hours. The cells were stained for spike protein and the percentage of cells expressing the spike protein in each condition was plotted. All cannabinoids tested were isolated from a hemp extract as described in Methods. (B) Chemical structures of cannabinoids and 7-OH CBD. (C) A549-ACE2 cells were treated with indicated doses of 7-OH CBD followed by infection with the SARS-CoV-2 at an MOI of 0.5. The cells were stained for spike protein and the percentage of cells expressing the spike protein in each condition was plotted. Representative data of CBD from Figure 1C (Supplier A) is used for comparison. EC50 values are indicated.

CBD is rapidly metabolized in the liver and gut into two main metabolites, 7-carboxy-cannabidiol (7-COOH-CBD) and 7-hydroxy-cannabidiol (7-OH-CBD). Although the levels of 7-COOH-CBD are 40-fold higher than 7-OH-CBD in human plasma, 7-OH-CBD is the active ingredient for the treatment of epilepsy (*14*). Like CBD but unlike the other cannabinoids, 7-OH-CBD effectively inhibited SARS-CoV-2 replication in A549-ACE2 cells (EC50 3.6 μM; Fig. 2C) and was non-toxic to cells (fig. S2H). Analysis of blood plasma levels in healthy patients taking FDA-approved CBD (Epidiolex^®^) shows a maximal concentration (C_max_) for CBD in the nM range whereas 7-OH-CBD had a C_max_ in the μM range, similar to that observed in cultured cells (*15*). These results suggest that CBD itself is not present at the levels needed to effectively inhibit SARS-CoV-2 in people. By contrast, the plasma concentrations of its metabolite 7-OH-CBD, whose C_max_ can be increased several-fold by co-administration of CBD with a high-fat meal, are sufficient to potentially inhibit SARS-CoV-2 infection in humans (*15*).

CBD could be acting to block viral entry to host cells or at later steps following infection. As CBD was shown to decrease ACE2 expression in some epithelial cells including A549 (*16*), we first determined whether CBD suppressed the SARS-CoV-2 receptor in our A549-ACE2 overexpressing cells. No decrease in ACE2 expression was observed (Fig. 3A). Furthermore, analysis of lentiviruses pseudotyped with either the SARS-CoV-2 spike protein or the VSV glycoprotein (*17*) showed that antibody to the spike protein effectively blocked viral infection of the SARS-CoV-2, but not VSV-G expressing viruses. However, 10 μM CBD only partially inhibited cell entry by spike-expressing virus, suggesting that other mechanisms were largely responsible for its antiviral effects (Fig. 3B, and figs. S3 A and B). By contrast, antibodies to the spike protein effectively blocked viral infection of the SARS-CoV-2 but not VSV-G expressing viruses. Consistent with this, CBD was also effective at inhibiting SARS-CoV-2 spike protein expression in host cells even 2 hours after infection in the presence of antibodies to the spike protein to prevent reinfection during this time period (Fig. 3C,D). To assess whether CBD might be preventing viral protein processing by the viral proteases Mpro or PLpro, we assayed their activity *in vitro* (fig. S4). CBD did not affect the activity of either protease, raising the possibility that CBD targets host cell processes.

**Fig. 3.**
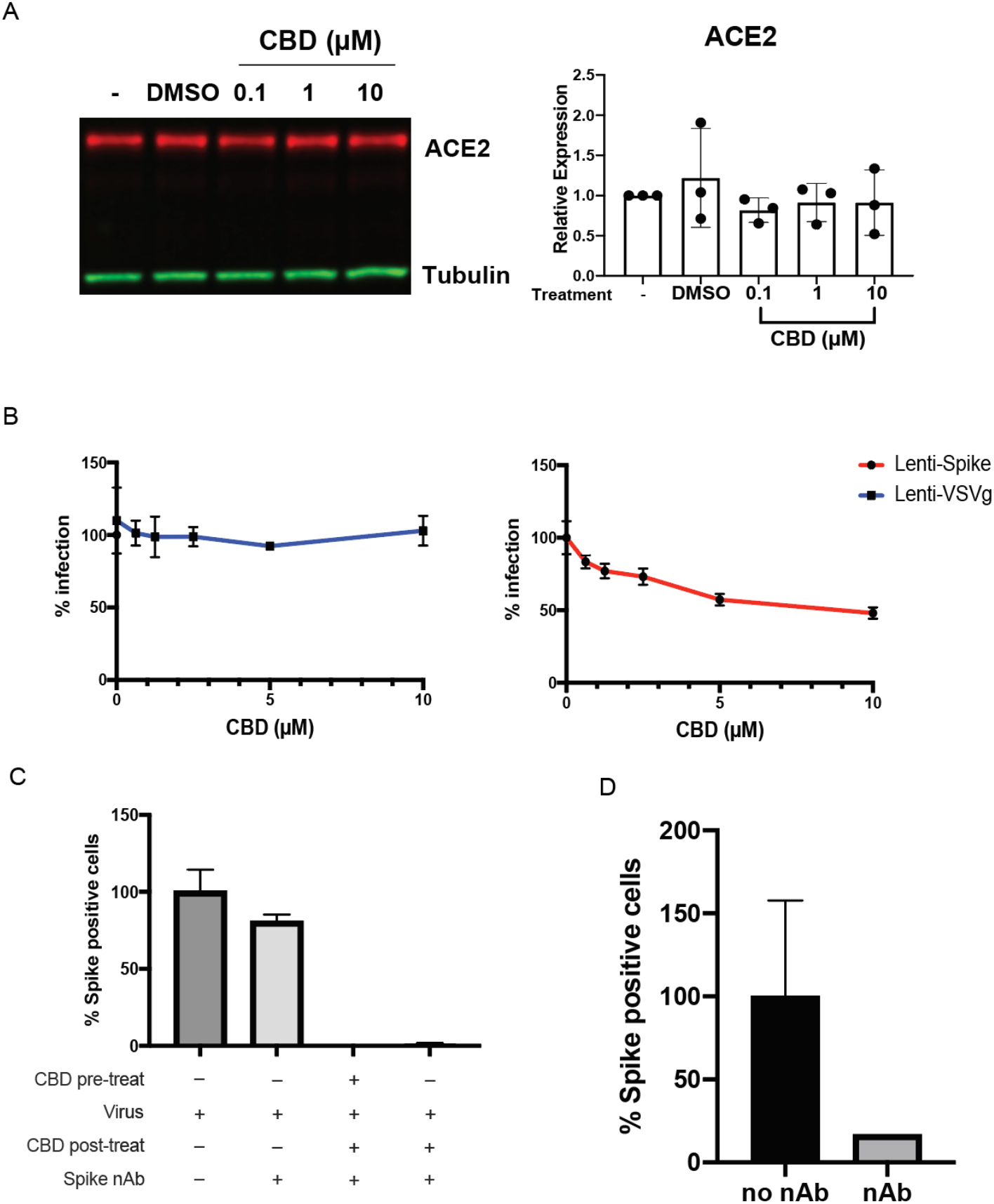
CBD inhibits viral replication after SARS-CoV-2 entry into the host cell. (A) Immunoblots of ACE2 protein expression from A549-ACE2 cell lysates either untreated or treated with vehicle or CBD at indicated doses (n=3). Blots were probed with antibodies against ACE2 and tubulin. ACE2 protein expression levels were normalized to the tubulin signal within each sample. ACE2 expression levels were plotted relative to untreated samples. Expression levels were compared to vehicle by one-way ANOVA. (B) 293T-ACE2 cells were infected by spike or VSV-G pseudovirus for 72 hours with the indicated doses of CBD treatment, and the percentage of infected cells plotted. (C) A549-ACE2 cells were either pre-treated or not with 10 μM CBD for 2 hours, then infected with SARS-CoV-2 at an MOI of 0.5 for 2 hours. Cells were then treated with 10 μM CBD or DMSO for 16 hours with the spike neutralizing antibody to prevent reinfection. Spike positive cells were quantified and normalized to the virus-infected only sample. (D) Validation of neutralizing antibody efficacy. 400 pfu of SARS-CoV-2 virus was incubated with or without 100 μM of neutralizing antibody for 1 hour. A549-ACE2 cells were treated with the mixture for 16 hours and Spike positive cells were quantified.

Consistent with this interpretation, RNA-seq analysis of infected A549-ACE2 cells treated with CBD for 24 hours shows a striking suppression of SARS-CoV-2-induced changes in gene expression. CBD effectively eradicated viral RNA expression in the host cells, including RNA coding for spike, membrane, envelope and nucleocapsid proteins (Figs. 4 A and B). Both SARS-CoV-2 and CBD each induced significant changes in cellular gene expression, including a number of transcription factors (figs. S5 and S6). Principal component analysis of host cell RNA shows almost complete reversal of viral changes but, rather than returning to a normal cell state, the CBD+virus infected cells resemble those treated with CBD alone (Fig. 4C). Clustering analysis using Metascape reveals some interesting patterns and associated themes (Fig. 4D, figs. S7, and S8). For example, viral induction of genes associated with chromatin modification and transcription (Cluster 1) is reversed by CBD although CBD alone has no effect. Similarly, viral inhibition of genes associated with ribosomes and neutrophils (Cluster 3) is largely reversed by CBD, but the drug alone has no effect. This contrasts with Clusters 5 and 6 where CBD alone induces strong activation of genes associated with the host stress response. Together these results suggest that CBD acts to prevent viral protein translation and associated cellular changes.

**Fig. 4.**
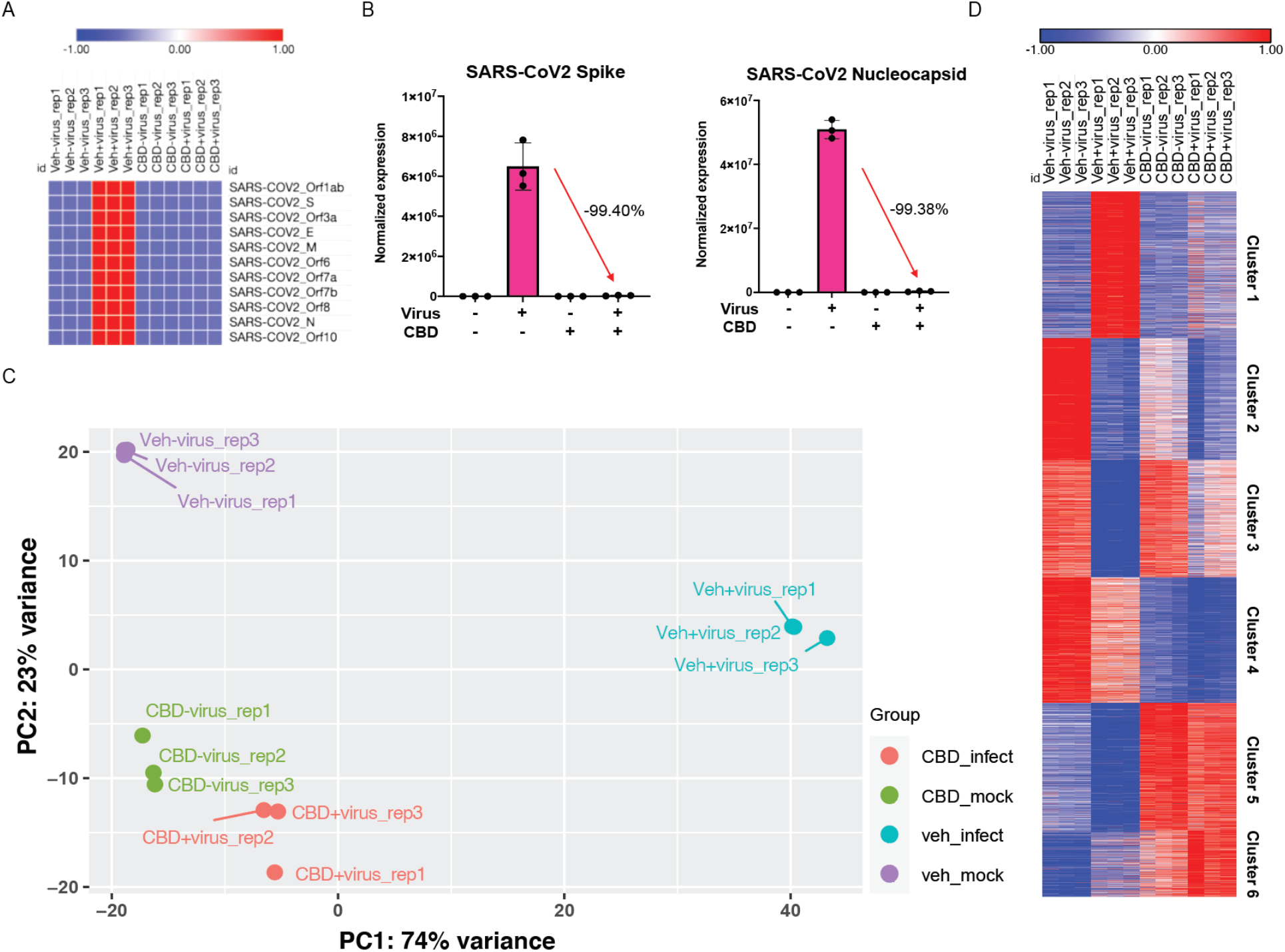
Changes in Viral and host cell transcription following SARS-CoV-2 infection or CBD treatment. A549-ACE2 cells were infected with SARS-CoV-2 at MOI of 3 with or without CBD treatment at 10 μM for 24 hours. RNA-seq was performed as described in Methods. (A) Heatmap of relative levels of SARS-CoV-2 genes from the RNA-seq samples. (B) Expression levels of SARS-CoV-2 spike and nucleocapsid genes. Percent expression level changes for genes from infected cells compared to cells infected and CBD treated are indicated for each gene. (C) Principal component analysis (PCA) of RNA-seq data showing control (veh_mock), SARS-CoV-2 infected (veh_infect), CBD-treated (CBD_mock), and SARS-CoV-2 infected plus CBD treated (CBD_infect) samples. The first and second principal components (PC1 and PC2) of each sample are plotted. (D) Heatmap of normalized expression levels of 5,000 most variable genes across all RNA-seq samples, clustered into 6 groups based on differential expression between treatment conditions.

One potential mechanism by which CBD could suppress viral infection and promote degradation of viral RNA is through induction of the interferon signaling pathway. Interferons are among the earliest innate immune host responses to pathogen exposure (*18*). SARS-CoV-2 infection suppresses the interferon signaling pathway (*19*) (Fig. 5A, and fig. S9). Some genes that are induced by CBD in both the absence and presence of the virus include receptors for interferons beta and gamma as well as mediators of the signaling pathway such as STATs 1 and 2 (Fig. 5A and fig. S10). Other genes in the pathway like OAS1, an interferon-induced gene that leads to activation of RNase L and RNA degradation (*20*), are not significantly induced by CBD unless in the presence of the virus (Fig. 5A and fig. S11). These latter results are consistent with the possibility that CBD lowers the effective viral titer sufficiently to enable normal host activation of the interferon pathway. At the same time, CBD effectively reverses viral induction of cytokines that can lead to the deadly cytokine storm at later stages of infection (Fig. 5B). Collectively, these results suggest that CBD inhibits SARS-CoV-2 infection in part by activating the interferon pathway leading to degradation of viral RNA and subsequent viral-induced changes in host gene expression, including cytokines.

**Fig. 5.**
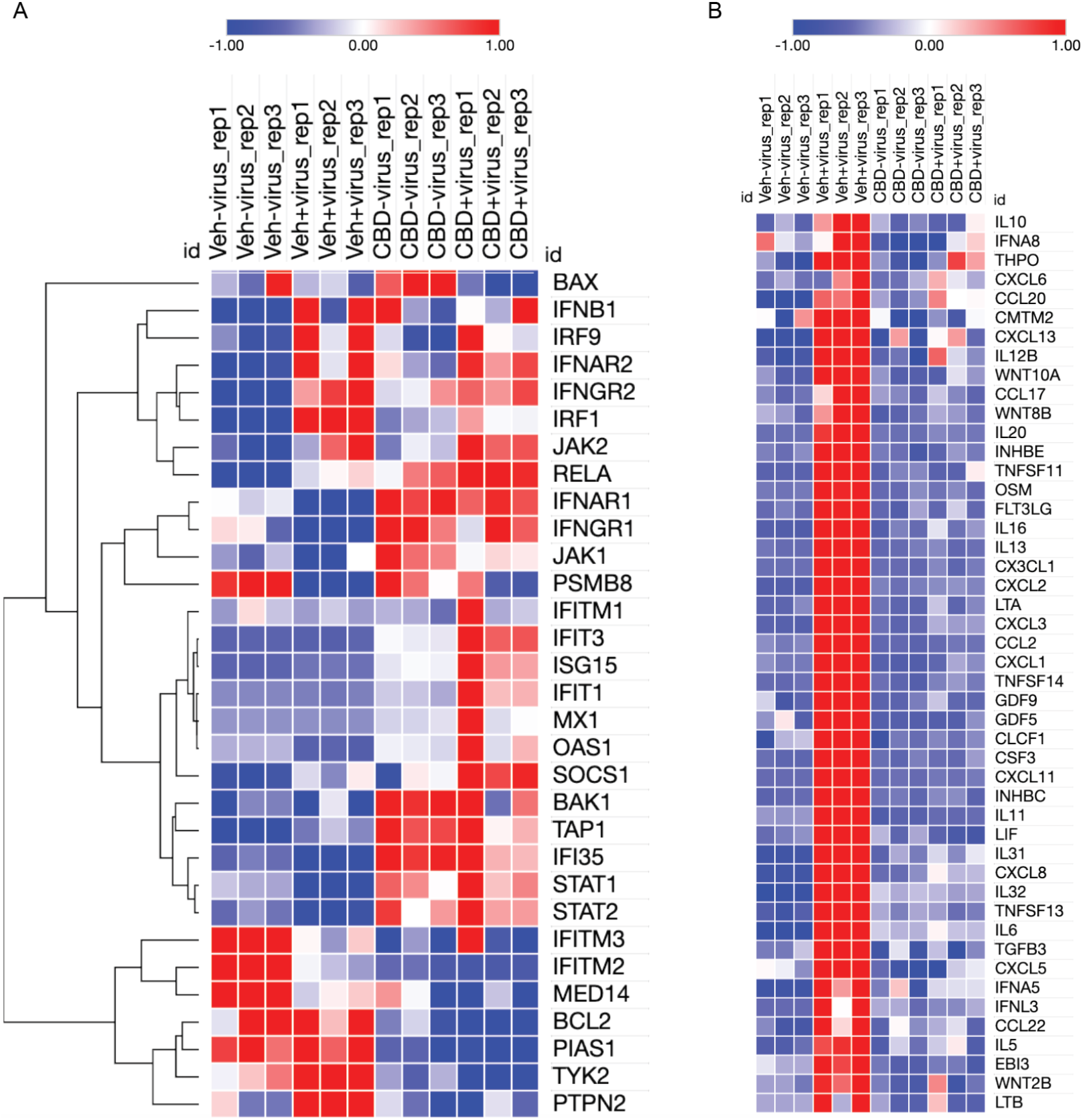
CBD promotes host cell interferon responses and inhibits viral induction of cytokines. (A) Heatmap of normalized expression levels of genes from the Interferon Response Canonical Pathway for all RNA-seq samples including control (veh_mock), SARS-CoV-2 infected (veh_infect), CBD-treated (CBD_mock), and SARS-CoV-2 infected plus CBD treated (CBD_infect) samples. Hierarchical clustering was applied to the genes. (B) Heatmap of normalized expression levels of GO Cytokine Activity genes which were up-regulated by the viral infection but down-regulated by CBD treatment for all RNA-seq samples as described in (A).

Given that CBD preparations containing substantial amounts of CBD are taken by a large number of individuals, we examined whether CBD exposure might correlate to a decreased risk of SARS-CoV-2 infection. Analysis of over 93,000 patients tested for SARS-CoV-2 at the University of Chicago Medical Center showed that 10.0% tested positive overall, but only 5.7% of the ~400 who had any cannabinoid in their medical record tested positive (Fig. 6). Patients taking CBD versus other cannabinoids had an even lower rate of testing positive (1.2% in 85 CBD patients versus 7.1% in 113 patients taking other cannabinoids, p=0.08). This finding that patients taking other cannabinoids had less protection against viral infection is consistent with our cell culture studies. Since multiple potential confounding factors could explain these findings, including age, race, clinical morbidities, and sex, we matched 82 patients who were prescribed oral, FDA-approved CBD (Epidiolex^®^) before COVID-19 testing to patients who had no indication of taking any cannabinoids but had comparable other characteristics including similar demographic characteristics, clinical comorbidities, and records of other medications in the two years before COVID-19 testing (table S1). Of the patients prescribed oral CBD before their COVID-19 test, the most common morbidity categories were hypertension and conditions with immunosuppression. Strikingly, only 1.2% of the patients prescribed CBD contracted SARS-CoV-2 whereas 12.2% of the matched, non-cannabinoid patients tested positive (p=0.009), suggesting a potential reduction in SARS-CoV-2 infection risk of approximately an order of magnitude.

**Fig. 6.**
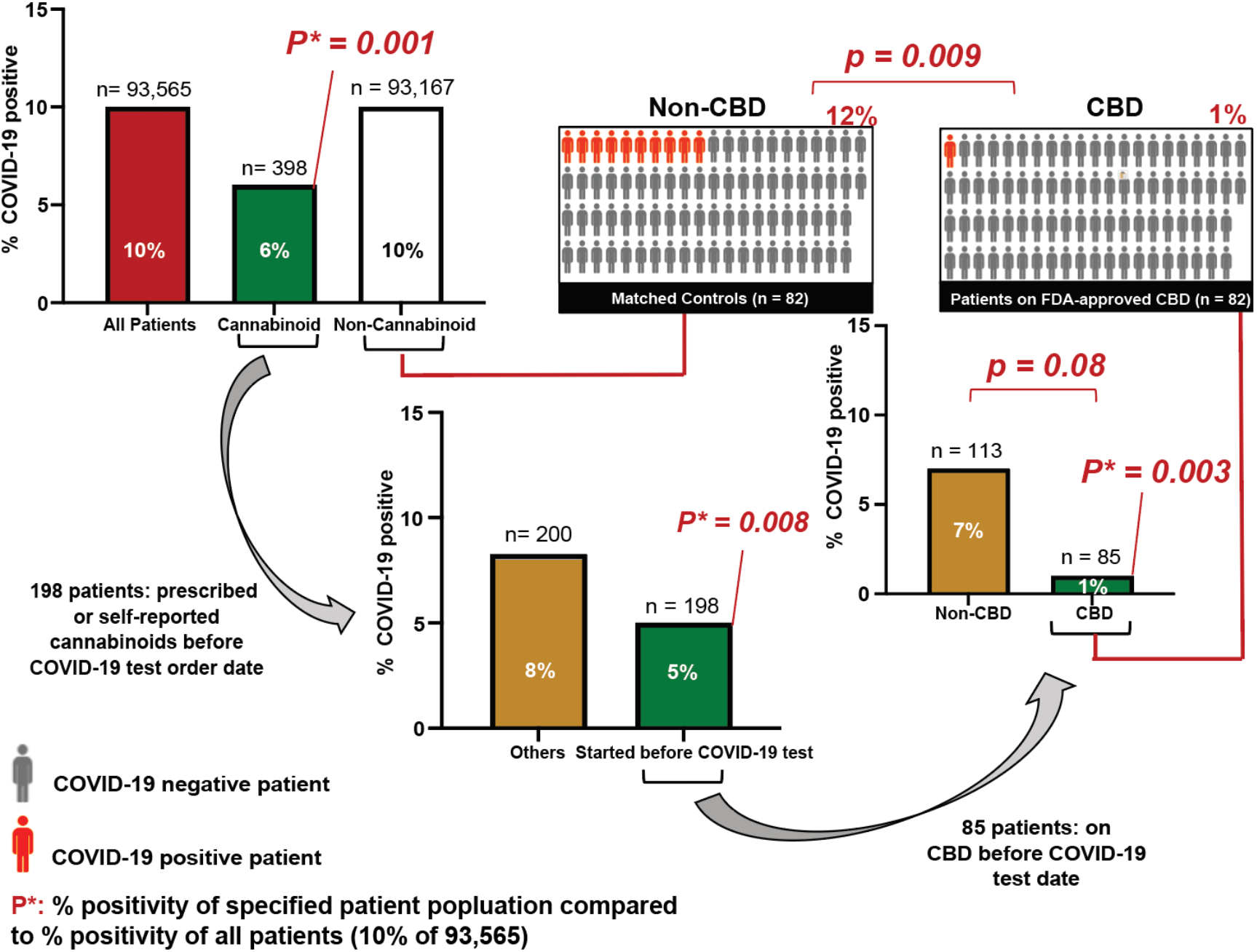
High Dose CBD usage in patients is significantly correlated with a reduction in COVID-19 positivity. Associations between reported cannabinoid medication use and COVID-19 test results among adults tested at the University of Chicago Medicine (total n=93,565). P*: p-values of percent positivity of the specified patient population compared to percent positivity of all patients (10% COVID-19 positive among 93,565 patients). Middle right: 85 patients took CBD before their COVID test date. Upper right: 82 of the 85 patients took FDA-approved CBD (Epidiolex^®^) and were matched to 82 of the 93,167 patients (Matched Controls) with a nearest neighbor propensity score model that scored patients according to their demographics and their recorded diagnoses and medications from the two years before their COVID-19 test. P-values were calculated using Fisher’s exact test two-sided.

## DISCUSSION

Our results suggest that CBD can block SARS-CoV-2 infection at early stages of infection, and CBD administration is associated with a lower risk of SARS-CoV-2 infection in humans. Furthermore, the active compound in patients is likely to be 7-OH-CBD, the same metabolite implicated in CBD treatment of epilepsy. The substantial reduction in SARS-CoV-2 infection risk of approximately an order of magnitude in patients who took FDA-approved CBD highlights the potential efficacy of this drug in combating SARS-CoV-2 infection. Finally, the ability of CBD to inhibit replication of MHV raises the possibility that CBD may have efficacy against new pathogenic viruses arising in the future.

One mechanism contributing to the antiviral activity of CBD is the induction of the interferon pathway both directly and indirectly following activation of the host immune response to the viral pathogen. In fact, interferons have been tested clinically as potential treatments for COVID-19 (*21*). Importantly, CBD also suppresses cytokine activation in response to viral infection, reducing the likelihood of immune cell recruitment and subsequent cytokine storms within the lungs and other affected tissues. These results complement previous findings suggesting that CBD suppresses cytokine production in recruited immune cells such as macrophages (*22*). Thus, CBD has to the potential not only to act as an antiviral agent at early stages of infection but also to protect the host against an overactive immune system at later stages.

CBD has a number of advantages as a potential preventative agent against SARS-CoV-2. CBD is widely available without restricted access if the content of THC is <0.3%. There are multiple means of ingestion, including potential for inhalation and nasal delivery. CBD blocks viral replication after entry into cells and, thus, is likely to be effective against viral variants with mutant spike proteins. Unlike drugs such as remdesivir or antiviral antibodies, CBD administration does not require injection in hospital settings. Finally, CBD is associated with only minor side effects (*15*).

However, several issues require close examination before CBD can be considered or even explored as a therapeutic for COVID-19 (*11*). Although many CBD formulations are available on the market, they vary vastly in quality, the amount of CBD, and their pharmacokinetic properties after oral administration, which are mostly unknown. CBD is quite hydrophobic and forms large micellar structures that are trapped and broken down in the liver, thereby limiting the amount of drug available to other tissues after oral administration. The inactive carriers have a significant impact on clinically obtainable concentrations. As CBD is widely sold as a preparation in an edible oil, we analyzed flavored commercial hemp oils and found a CBD content of only 0.30% in a representative sample (fig. S12). The purity of CBD and, in particular, the composition of the materials labelled as CBD are also important, especially in light of our findings suggesting that other cannabinoids such as THC might act to counter CBD antiviral efficacy. This essentially eliminates the feasibility of marijuana serving as an effective source of antiviral CBD, in addition to issues related to its legal status. Finally, other means of CBD administration such as vaping and smoking raise concerns about potential lung damage.

Future studies to explore the optimal means of CBD delivery to patients along with clinical trials will be needed to fully test the promise of CBD as a therapeutic to block SARS-CoV-2 infection. As the clearance rates for CBD in plasma are substantially lower in humans than mice, we would suggest moving to clinical trials rather than doing preclinical studies in animal models (*15*). We advocate carefully designed placebo-controlled clinical trials with known concentrations and highly-characterized formulations in order to define CBD’s role in preventing and treating early SARS-CoV-2 infection. The necessary human *in vivo* concentration and optimal route and formulation remain to be defined. We strongly caution against the urge to take CBD in presently available formulations as a preventative or treatment therapy at this time, especially without the knowledge of a rigorous randomized clinical trial with this natural product (*23*).

## ACKNOWLEDGEMENTS

We thank the members of the SARS-CoV-2 host response team in Chicago for stimulating discussions and support with particular thanks to Julian Solway, Rick Morimoto, Nissim Hay, Anne Sperling, HuanHuan Chen, Raphael Lee, Raymond Roos, Shannon Elf, Alexander Muir, Gokhan Mutlu, Jay Pinto, Steven White, Nickolai Dulin, Ray Moellering, Viswanathan Natarajan, Leonitis Platanias, Karen Ridge and HuanHuan Chen. We thank Dominique Missiakas for facilitating access to the University of Chicago Howard Taylor Ricketts Facility by providing protocols and trained scientists. We also thank Nicole Rosner and Kathleen Cagney for proposing and facilitating analysis of clinical data, and Mark Ratain for consideration of pharmacokinetic issues. We thank the University of Chicago Genomics Facility (RRID:SCR_019196) especially Sandhiya Arun and Pieter Faber, for their assistance with RNA sequencing. Finally, we would like to acknowledge the University of Chicago Vice Provost for Research, Karen Kim, and the Dean of the Biological Sciences Division, Kenneth Polonsky, for their steadfast support.

## Author contributions

Conceptualization: MRR, LCN, DY, GR, SAO, GFP, DOM, ND

Methodology: MRR. GR, SAO, DOM, GFP, ND, BCD

Software: DY, TB

Formal analysis: DY, TB

Investigation: LCN, DY, VN, TB, TO, SC, JBF, ND, AM, CD, DS, HG, KAJ

Resources: GR, MRR, GFP, ST, BCD

Data Curation: DOM, TB, DY

Writing – original draft: MRR

Writing – review & editing: LCN, DY, VN, TB, SC, JBF, ND, AM, KAJ, DS, JMM, BCD, ST, SAO, GFP, DOM, GR, MRR

Visualization: LCN, DY, ND, AM, SC, BCD, JBF

Supervision: MRR. GR, DOM, GFP, SAO, ST, BCD

Project Administration: MRR

Funding acquisition: MRR

## Data and materials availability

All data, code, and materials used in the analysis will be available in some form to any researcher for purposes of reproducing or extending the analysis except when limited by materials transfer agreements (MTAs). Raw and processed RNA-seq data will be deposited into the GEO database.

## Supplementary Materials

Materials and Methods

Figs. S1 to S12

Table S1

