## Supplementary Materials for "Cannabidiol Inhibits SARS-CoV-2 Replication and Promotes the Host Innate Immune Response"

**This PDF file includes:**

Materials and Methods
Figs. S1 to S12
Table S1

### Materials and Methods

Materials and Cells

High-purity CBD and 7-OH-CBD were acquired from two chemical companies or an online commercial source. Cannabinoid-infused hemp oil containing 1,500+ mg cannabinoids was from Bluebird Botanicals (Louisville, CO, USA). Hemp extract from *C. sativa* biomass was from Hopsteiner Ltd. (Yakima, Washington, USA). Low CBD hemp oil was obtained from an online commercial source. KPT-9274 is from Selleckchem (Houston, TX). URMC-099 is from ApexBio (Houston, TX). A549-ACE2 cells were generously provided by tenOever and colleagues (*17*). Vero E6 cells were purchased from ATCC.

SARS-CoV-2 infection assay

All SARS-CoV-2 infections were performed in biosafety level 3 conditions at the Howard T. Ricketts Regional Biocontainment Laboratory. Cells in DMEM +2% FBS were treated with CBD or other inhibitors or 2 hours with 2-fold dilutions beginning at 10 μM in triplicate for each assay. A549-ACE2 cells were infected with an MOI (multiplicity of infection) of 0.5 in media containing the appropriate concentration of drugs. Vero E6 cells were infected with an MOI of 0.1 in media containing the appropriate concentration of drugs. After 48 hours, the cells were fixed with 3.7% formalin, blocked, and probed with mouse anti-Spike antibody (GTX632604, GeneTex) diluted 1:1,000 for 4 hours, rinsed, and probed with anti-mouse-HRP for 1 hour, washed, then developed with DAB substrate 10 minutes. Spike positive cells (n>40) were quantified by light microscopy as blinded samples. Data were analyzed and plotted using Prism and EC50 values were extracted from nonlinear fit of response curves.

Crystal Violet toxicity assay

Cells were treated with varying concentrations of different compounds in 2% DMEM starting at 10μM and going down by 1/2 for six more dilutions. Cells were incubated with the drug for 48 hours. Cells were fixed with 10% Formalin solution for 30 minutes. Then they were stained with 1% Crystal Violet solution for 30 minutes after which plates were dried and the amount of Crystal Violet staining was assessed by measuring absorbance at 595nm on a TECAN M200 Plate reader. Absorbance readings were normalized to those of the control wells not treated by the drug to measure the differences in cell growth with or without the drug treatment.

Spike protein antibody neutralizing assay

A549-ACE2 cells were treated with 10 μM of CBD either 2 hours before infection or 2 hours after infection. Cells were infected with MOI of 3 for 2 hours. Then, the infection media was replaced with media containing CBD or DMSO, and 100 μM of neutralizing antibody (Active Motif 001414), and the samples were incubated at 37°C for 16 hours. After which the samples were fixed with 10% formalin and underwent IHC for spike protein. Neutralizing antibody efficiency was tested by incubating 400 pfu of virus with or without 100 μM of the antibody at 37°C for 1 hour. Then A549-ACE2 cells were infected with the mixture for 16 hours. Spike positive cells were quantified as described above.

MHV infection assay

Recombinant MHV-A59 virus containing a Fluc reporter gene was obtained from the Gallagher lab at Loyola's Stritch School of Medicine(*22*). For firefly luciferase reporter assay, A549-MHVR cells were seeded in a 96-well plate with 5,000 cells per well and pre-treated for 2 hours with the following treatments: remdesivir (MedChemExpress) and CBD (Cayman). After 2 hours, MHV-A59 virus was added at an MOI of 0.1 for 12 hours. At the end of infection, cells were washed with 1 x PBS and then lysed using 1x lysis buffer (Promega) and placed on a shaker for 10 minutes. 20 µL of lysis was then added to a black 96-well black, flat bottom plate (Costar) where 50 µL of 1x homemade luciferase substrate was previously added (1 mM D-luciferin, 3 mM ATP, 15 mM MgSO4·H2O, 30 mM HEPES [pH 7.8]) and using a BioTek microplate luminometer the plate was read for luminescence. For assessing toxicity, 50,000 A549-MHVR cells were seeded in a 24-well plate. Cells were transfected with renilla plasmid DNA for 24 hours using Lipofectamine 2000 (Invitrogen) according to the manufacturer's instructions. After a 24-hour transfection, cells were treated with remdesivir or CBD for 12 hours. At the end of treatment, cells were washed with 1x PBS and then lysed using 1x lysis buffer (Promega) and placed on a shaker for 10 minutes. 20µl of lysis buffer was then added to a black 96-well flat bottom plate (Costar) where 50 µl of renilla buffer (*23*) was previously added. The plate was read for luminescence using a BioTek microplate luminometer.

Description of the Cannabinoids

CBD can be procured by isolating cannabidiolic acid (CBDA) from *Cannabis sativa* plant material and then inducing chemical decarboxylation, or via decarboxylation of raw plant material or extract and subsequent isolation of CBD. Cannabidivarin (CBDV) is a naturally occurring CBD homolog that has an *n*-propyl in place of CBD's *n*-pentyl side chain. Cannabigerol (CBG), in the form of cannabigerolic acid, is the metabolic precursor to both tetrahydrocannabidiolic acid and CBDA in *C. sativa*. Tetrahydrocannabinol (THC) is a cyclized congener of CBD that is obtained from tetrahydrocannabinolic acid decarboxylation. THC is present in *C. sativa* in both $\Delta^9$-*cis* and $\Delta^9$-*trans* stereoisomers. Cannabichromene (CBC), in the form of cannabichromenic acid, represents a third possible cannbigerolic acid metabolite.

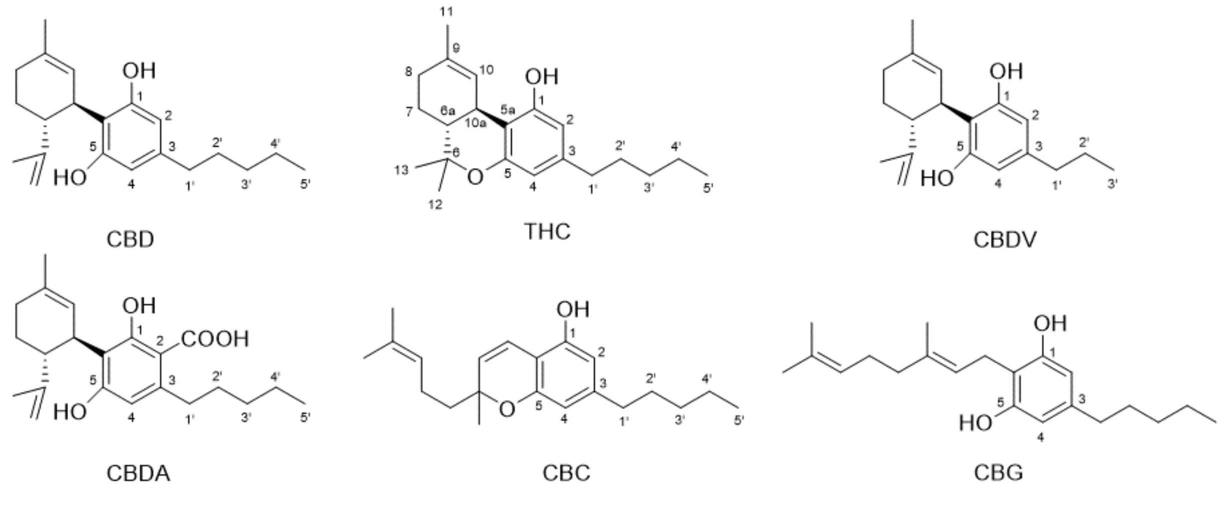


Acquisition, Isolation and Characterization of Cannabinoids

In the present study, purification of CBD from natural sources used (a) cannabinoid infused hemp oil containing 1,500+ mg cannabinoids in medium-chain triglycerides per fluid ounce, manufactured by Bluebird Botanicals (Louisville, CO, USA), and (b) hemp extract prepared by supercritical fluid extraction (SFE) with $CO_2$ from *C. sativa* biomass qualifying as hemp, manufactured by Hopsteiner Ltd. (Yakima, Washington, USA) with a 54.7% total content of CBD, calculated as CBD+CBDa*0.877. Typical purities of these CBD preparations are in the 90-97% range including foreign impurities (e.g., residual solvent). Details of the purification and structure analysis methodologies are detailed in a concurrent publication, which is currently under review elsewhere. In brief, the methodologies can be summarized as follows:

*Purification Procedure.* CBD, CBC, CBG, $\Delta^9$-*trans*-THC, $\Delta^9$-*cis*-THC, and CBDV were isolated from the hemp oil, and CBDA from the crude hemp SFE extract, using centrifugal partition chromatography (CPC), a countercurrent separation technique, and a biphasic liquid-liquid solvent system.

*Structure Elucidation Methodology*. The identity of the commercially sourced CBD samples was verified by 1D $^1$H NMR analysis, performed as qNMR measurement, via comparison with an authentic HiFSA profile of CBD as published (*21*). In addition to an overall excellent match of the profiles, the highly coupled fingerprint signal of H-4''ax served as a highly specific identity marker. The structures of the cannabinoids purified from the natural sources were established by a combination of 1D/2D NMR and LC-HRMS analysis, taking into account reference data from the literature.

*NMR Sample Preparation.* For commercial samples supplied as solution, the solvent was removed carefully in vacuo and 450μL deuterated methanol (MeOH-$d_4$) added to the residue using a precision syringe. The solution was transferred into 5-mm NMR tube with a glass pipette, the vial rinsed three times with 25μL of solvent and the rinsing solution transferred into the same NMR tube, for a final volume of 525μL. Commercial and isolated samples available as solids were directly weighed into a 5-mm NMR tube and 500μL of solvent added with a precision syringe. For analysis of the commercial hemp oil preparation, 10 drops (0.25 mL equivalent to 14-15 drops) was added into 5 mm NMR tube directly. The net weight of hemp oil in NMR tube is 198.50 mg on a 0.01 mg precision balance. 0.90 mg of dinitrobenzoic acid was serviced as an internal reference standard. 325μL and 10μL deuterated NMR chloroform and methanol were added, then the tube was flame sealed.

NMR Data Acquisition and Processing and qNMR Evaluation

All $^1$H NMR data were acquired on a Bruker 600 Avance III with a two channel $^{13}$C direct cryogenic probe. Time domain (TD) was set to 64k, relaxation delay (D1) was 60 sec, 90-degree excitation pulses were used for a total of 32 signal averaged scans. The receiver gain (RG) was 32 for all samples, except for one mass-limited sample <1mg (RG = 101) and the large-quantity hemp oil sample (RG = 2; 15 degrees excitation pulse used). Determination of sample purity and CBD content in hemp oil by quantitative NMR (qNMR) using the 100% qNMR approach and openly published worksheets (https://gfp.people.uic.edu/qnmr/content/qnmrcalculations/100p.html). The qNMR purity of all CBD samples was >97% including foreign impurities, and no cannabinoid congeners could be detected at levels of>1.0%. Using the absolute qHNMR method with internal calibration (IC abs-qNMR), the content of CBD in hemp oil was determined as 0.30%.

Pseudotyped Lentivirus Production

293T and 293T-ACE2 cells were cultured in DMEM (Corning 10017CV) with 1X sodium pyruvate (Gibco 11360070) and 10% FBS (HyClone SH30910). Lentivirus particles pseudotyped with SARS-CoV-2 (Wuhan-Hu-1) spike protein or VSV-G, were generated as described (*19*). Briefly, 293T cells were transfected using TransIT-LT1 (Mirus) with 3rd generation lentivirus packaging vectors (HDM-Hgpm2, HDM-tat1b, pRC-CMV-Rev1b), transfer vector (pHAGE-CMV-ZsGreen-W) and either SARS-CoV-2 spike (HDM-IDTSpike-fixK) or VSV-G (HDH-VSVG). Supernatants collected at 36 and 60 hours post-transfection were pooled, syringe filtered and frozen in single-use aliquots at -80°C. All plasmids used for lentivirus production were kindly provided by Dr. Jesse Bloom (University of Washington, Seattle).

Pseudovirus Binding Assay

293T-ACE2 cells were seeded at $1.2\times10^4$ cells per 96well in black wall, clear bottom plates. The next day, 2-fold dilutions of CBD stock (10mM) were prepared in DMSO, followed by 1:1000 dilutions in either complete DMEM or pseudovirus preparation. SARS-CoV-2 spike pseudovirus was used undiluted, while VSV-G pseudovirus was diluted 1:1,500 in complete DMEM. Cells and pseudovirus were pre-treated with CBD dilutions for 2 hours and 1 hour at 37°C, respectively. Cells were infected with pseudovirus for 72 hours, fixed with 4% paraformaldehyde, stained with a nuclear marker (Hoechst 33342, ThermoFisher H3570) and imaged. 293T-ACE2 cells were generously supplied by Dr. Jesse Bloom (University of Washington, Seattle).

Pseudovirus neutralization assay

293T or 293T-ACE2 cells were seeded at $1.2\times10^4$ cells per 96well in black wall, clear bottom plates. Next day, SARS-CoV-2 spike neutralizing antibody (Sino Biological 40592-R001) was diluted in complete DMEM to a starting final concentration of 300ng per 100ul per 96well, followed by subsequent 3-fold dilutions. Neutralizing antibody was incubated with pseudovirus for 1hr at 37°C. Cells were infected with pseudovirus +/- neutralizing antibody for 72hrs, fixed with 4% paraformaldehyde, stained with nuclear marker Hoechst 33342 and imaged.

Protease inhibition assay

Assays were performed in duplicate at room temperature in 96-well black plates at 25 °C. Reactions containing varying concentrations of inhibitor (10 or 50 μM) and 3CLpro enzyme (0.4 μM) or PLpro enzyme (0.3 μM) in Tris-HCl pH 7.3, 1mm EDTA were incubated for approximately five minutes. 3CLpro reactions were then initiated with TVLQ-AMC probe substrate (40 μM) and PLpro reactions were initiated with LKGG-AMC probe substrate (40 μM). The reaction plate was shaken linearly for 5 s and then measured for fluorescence emission intensity (excitation λ: 364 nm; emission λ: 440 nm) over time (1 min-3 h) on a Synergy Neo2 Hybrid). Each assay contained 2-3 positive control wells (DMSO) and 2 negative control wells (assay components without protease). Data were normalized to the positive control wells at 3 h, which was assigned an arbitrary value of 100.

Immunoblotting

A549-ACE2 cells were treated with CBD, vehicle (DMSO) or not treated for 24 hours. Cells were first washed with ice-cold PBS. Whole-cell extraction were prepared by directly lysing cells with Laemmli sample buffer (Bio-rad 1610747) supplemented with protease inhibitor (Roche 4693159001), PMSF (Roche 10837091001) and phosphatase inhibitor (GB-450) at 4°C. Protein samples were finally boiled at 98°C for 5 mins. Western blotting was performed using antibodies for ACE2 (Abcam 108252) and $\alpha$-tubulin (Invitrogen MA1-19401) for control. Blots were imaged and quantified using Licor Odyssey Fc.

RNA sequencing

Lung aveolar A549 cells were stably overexpressed with human angiotensin converting enzyme 2 (ACE2) protein and seeded at 10,000 cells per well in a 96-well plate. Cannabidiol or vehicle were added together to the cells. Cannabidiol (Cayman Chemical, 90080) was dissolved in a 10mM stock solution with DMSO (Sigma-Aldrich, D2650-100mL). Final concentration of CBD was 10 μM. The virus stock was then removed and replaced with fresh 2% FBS DMEM media with drug. The cells were incubated for another 24 hours before total RNA extraction the NucleoSpin 96 RNA kit (Takarabio, 740709).

Three independent biological replicates were performed per experimental condition, with 12 total RNA samples. RNA sample quality check, library construction, and sequencing were performed by the University of Chicago Genomics Facility following standard protocols. The average RNA Integrity Score was 8.9. All 12 samples were sequenced in two runs by a NovaSeq 6000 sequencer to generate paired-end 100bp reads. For each sample, the raw FASTQ files from two flow cells were combined before downstream processing.

RNA-seq data were analyzed using a local Galaxy 20.05 instance for the following steps (*24*). Quality and adapter trimming were performed on the raw sequencing reads using Trim Galore! 0.6.3 (*25*). The reads were mapped to both the human genome (UCSC hg19 with GENCODE annotation) and the SARS-COV-2 genome (NCBI Assembly ASM985889v3 with Ensembl annotation) using RNA STAR 2.7.5b (*26*). The resulting mapped reads from each sample were counted by featureCounts 1.6.4 (*27*) for per gene read counts. The raw counts were analyzed for differential expression between experimental conditions using DESeq2 1.22.1 (*28*), which also generated a normalized gene expression matrix and a PCA plot of the samples.

Clustering of variable genes

The top 5,000 most variable genes were selected, and the normalized gene expression data were analyzed by the Morpheus software (https://software.broadinstitute.org/morpheus). K-means clustering with 6 clusters was applied to the gene expression. For each gene, the normalized expression values of all samples were transformed by subtracting the mean and dividing by the standard deviation. The transformed gene expression values were used to generate the heatmap.

Interferon response

Expression data ($\log_2$ fold-change) and predicted activation status of genes were overlayed onto the interferon signaling pathway map using Ingenuity Pathway Analysis. Figures were generated through the use of IPA (QIAGEN Inc). Genes belonging to the

Ingenuity's interferon response Canonical Pathway were used for this heatmap. Normalized gene expression values were analyzed by the Morpheus software. Hierarchical clustering was performed on the genes using one minus Pearson correlation. For each gene, the normalized expression values of all samples were transformed by subtracting the mean and dividing by the standard deviation. The transformed gene expression values were used to generate the heatmap.

Gene set enrichment analyses

To identify themes across the 6 clusters, functional gene set enrichment analyses for the genes in each cluster were performed using Metascape (*29*). The following categories were selected for the enrichment analyses: GO Molecular Functions, KEGG Functional Sets, GO Biological Processes, Canonical Pathways, and KEGG Pathway. Additional parameters for Metascape: Min Overlap = 3, p-value Cutoff = 0.05, Min Enrichment = 1.5.

To identify gene sets which activities were reversed by CBD with viral infection, the input gene list includes genes significantly down-regulated by the virus (differential expression comparing veh-infect vs veh-mock, q-value cutoff 0.01) while also significantly up-regulated by CBD (differential expression comparing CBD-infect vs veh-infect, q-value cutoff 0.01). A second list includes genes significantly up-regulated by the virus (differential expression comparing veh-infect vs veh-mock) while also significantly down-regulated by CBD (differential expression comparing CBD-infect vs veh-infect). Gene set enrichment analyses were performed on these two lists of genes using the same Metascape method. Additionally, gene sets from the TRRUST database (*30*) were used to identify possible transcription factors regulating the genes.

Analysis of Patient Data

All patient data analysis was approved by the University of Chicago Biological Sciences Division institutional review board (IRB20-0842), which granted a waiver of consent because the data were deidentified except for elements of dates and because the risk to the privacy of participants was determined to be minimal.

*Participants:* Data was extracted from the University of Chicago Medicine (UCM) electronic medical record (EMR) for all UCM patients with a resulted COVID-19 polymerase chain reaction (PCR) test order date between March 3, 2020 and January 19, 2021 who were at least 18 years old on the test order date. To avoid potential confounding from intra-patients variability across multiple negative and/or positive tests during the time window of the study, we used a patient's first COVID-19 test order date in the data as the point of reference for classifying them either as COVID-19 Positive or COVID-19 Negative proximal to that date. In specific, a patient with a positive COVID-19 PCR test within 0 to 14 days on or after their first COVID-19 test order date was classified as Positive, otherwise, if the patient had only negative COVID-19 tests within those 14 days, they were classified as Negative.

*Measurements:*

*Medications.* All variables were obtained from the UCM EMR, Epic® (Epic Systems). COVID-19 status was determined by Centers for Disease Control or Viacor PCR tests until in-house testing with the Cobas® SARS-CoV-2 RT-PCR (Roche) began on March 15, 2020. Cannabinoid medication records were identified by searching the names of all provider-ordered and patient-reported medications documented in the UCM

EMR for the following substrings in either upper- or lower-case: *canemes, cannabi, cannibi, cbd, cesamet, dexanabinol, dronabinol, epidiolex, hemp, marijuana, marinol, nabilone, nabiximol, sativex, syndros, thc*. Only medications that had no end date or an end date between two years prior to their first COVID-19 test and January 19, 2021 were included in the search. Cannabinoids started before the date of the patient's first COVID-19 test order date were analyzed, and a cannabinoid was labeled Epidiolex® if its medication name contained the substring *epidiolex*. A patient was included in the Epidiolex Patients group if they had an Epidiolex® medication documented with a start date before their first COVID-19 test order date and had complete data on all demographic measurements included in Table S1. Non-cannabinoid medications were categorized according to the first six characters of their medication name in the UCM EMR.

*Morbidities*. Morbidity indicators were set equal to 1 if the patient had at least one of a list of ICD-10-CM diagnosis codes included on an administrative/billing record in the UCM EMR with a discharge date from 15 to 730 days before the patient's first COVID-19 test order. Morbidity diagnoses and medications started within 14 days before the patient's first COVID-19 test order were excluded to avoid confounding by potential early manifestations of COVID-19, e.g., presenting for care with symptoms that could lead to the diagnosis or medication. The lists of codes used for the morbidity indicators was defined using the HCUP Elixhauser Comorbidity Software (Agency for Healthcare Research and Quality). For each condition, all ICD-10-CM codes were used that are listed within categories in the Elixhauser file named Comorb_ICD10CM_Format_v2021-1.sas, which is entitled "Creation of Format Library for Comorbidity Groups, ICD-10-CM Comorbidity Software, Version 2021.1." The specific ICD-10-CM codes can be found by finding the following category variable names (in all caps) in the SAS file named above. Comorbidities with Immunosuppression: AIDS, ARTH, CANCER_LEUK, CANCER_LYMPH, CANCER_METS, CANCER_SOLID. Hypertension: CHFHTN_CX, CHFHTN_CXRENLFL_SEV, HTN_CX, HTN_CXRENLFL_SEV, HTN_UNCX. Chronic pulmonary disease: LUNG_CHRONIC. Depression: DEPRESS. Neurologic, seizures: NEURO_SEIZ. Liver disease: ALCOHOLLIVER_MLD, LIVER_MLD, LIVER_SEV. Neurologic, other: NEURO_OTH. Diabetes: DIAB_CX, DIAB_UNCX. Renal failure: CHFHTN_CXRENLFL_SEV, HTN_CXRENLFL_SEV, RENLFL_MOD, RENLFL_SEV. Neurologic, movement: NEURO_MOVT. Psychoses: DRUG_ABUSEPSYCHOSES, PSYCHOSES. Pulmonary circulation disorders: PULMCIRC. Dementia: DEMENTIA.

*Demographics*. Per UCM policy, sex, race, ethnicity, and other demographics are to be self-reported by patients by selecting from a pre-defined list of categories in the UCM EMR during patient registration. In some circumstances, nurses or other care providers will select demographics for patients in consultation with others whenever possible; for example, a nurse will occasionally select a patient's demographics based on consultation of available documentation or a patient's family members when the patient is unable to make their own demographic selections directly.

*Statistical analysis*. Epidiolex Patients were matched with patients with no record of any cannabinoid use using 1-to-1 nearest neighbor propensity score matching. The propensity scores used for matching were the estimated probabilities of taking Epidiolex® that were calculated from a multivariable logistic regression model that was fit to Epidiolex Patients and feasible controls. The probabilities were calculated using the *logit* then *predict* commands of Stata statistical software (Stata Corp). The covariates submitted to the propensity scoring model are listed in Table S1 along with their categorical frequencies or sample averages among Epidiolex Patients and among matched controls, which are compared using chi-squared tests or t-tests, respectively. Categorical characteristics were included as factor variables in the logit model and continuous measures were included as-is, except for age, which was included as both a continuous measure and a categorical measure. Controls were only considered feasible for matching if they had at least one medication record in the UCM EMR and had complete data available on all demographics used for matching. The nearest neighbor matching algorithm is described in Rassen et al. (*31*). It matched each Epidiolex Patient to a feasible control with the smallest (absolute) difference between its propensity score and the propensity score of the Epidiolex Patient. If a particular feasible control was closest to more than one Epidiolex Patient, the algorithm chose to match that control to the Epidiolex Patient that was the closest to that control and found different controls to match with the other Epidiolex Patients.

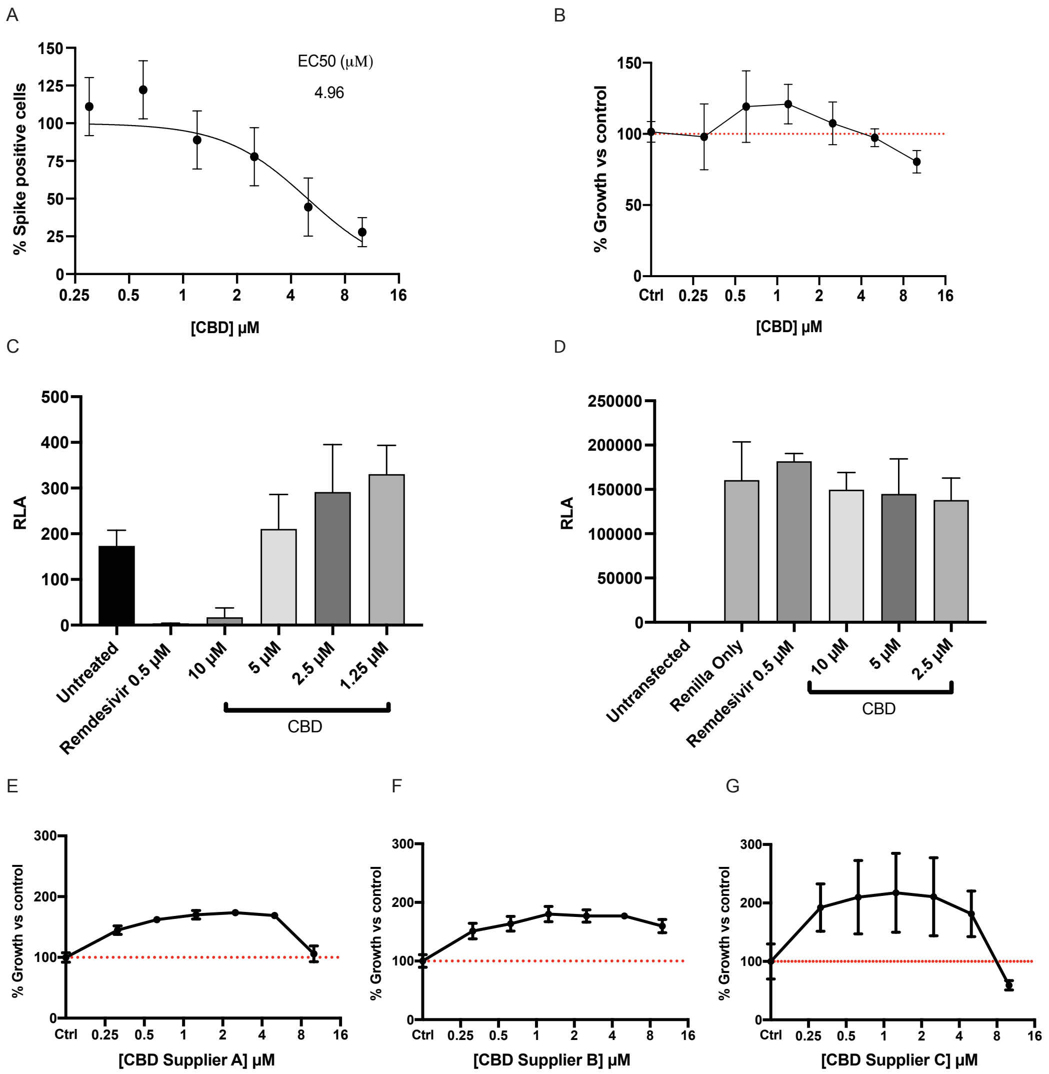


**Fig. S1. CBD inhibits SARS-CoV-2 and MHV replication in different cell lines *in vitro*.** (A) Vero E6 cells were treated with indicated doses of CBD followed by infection with SARS-CoV-2 at an MOI of 0.1 for 48 hours. The cells were stained for spike protein and the percentage of cells expressing the spike protein in each condition was plotted. The EC50 value is indicated. (B) Vero E6 cells were treated with indicated doses of CBD for 48 hours. The cells were stained by Crystal Violet as described in Methods and viability levels were plotted as cell growth relative to untreated controls. (C) A549-MHVR cells were infected with MHV-A59 at an MOI of 0.1 in the presence of decreasing concentrations of CBD and 0.5uM remdesivir. Virus entry was quantified by measuring relative luciferase activity (RLA) at 12 hours post-infection. (D) Cell viability of A549-MHVR cells treated with CBD was determined by transfecting cells with renilla plasmid DNA for 24 hours and then treating them with decreasing concentrations of CBD and 0.5uM remdesivir for 12 hours. Cell viability was quantified by measuring RLA at 12 hours post-infection. (E-G) A549-ACE2 cells were treated with indicated doses of CBD from three different suppliers A, B or C for 48 hours. Viability levels were plotted as described in (B).

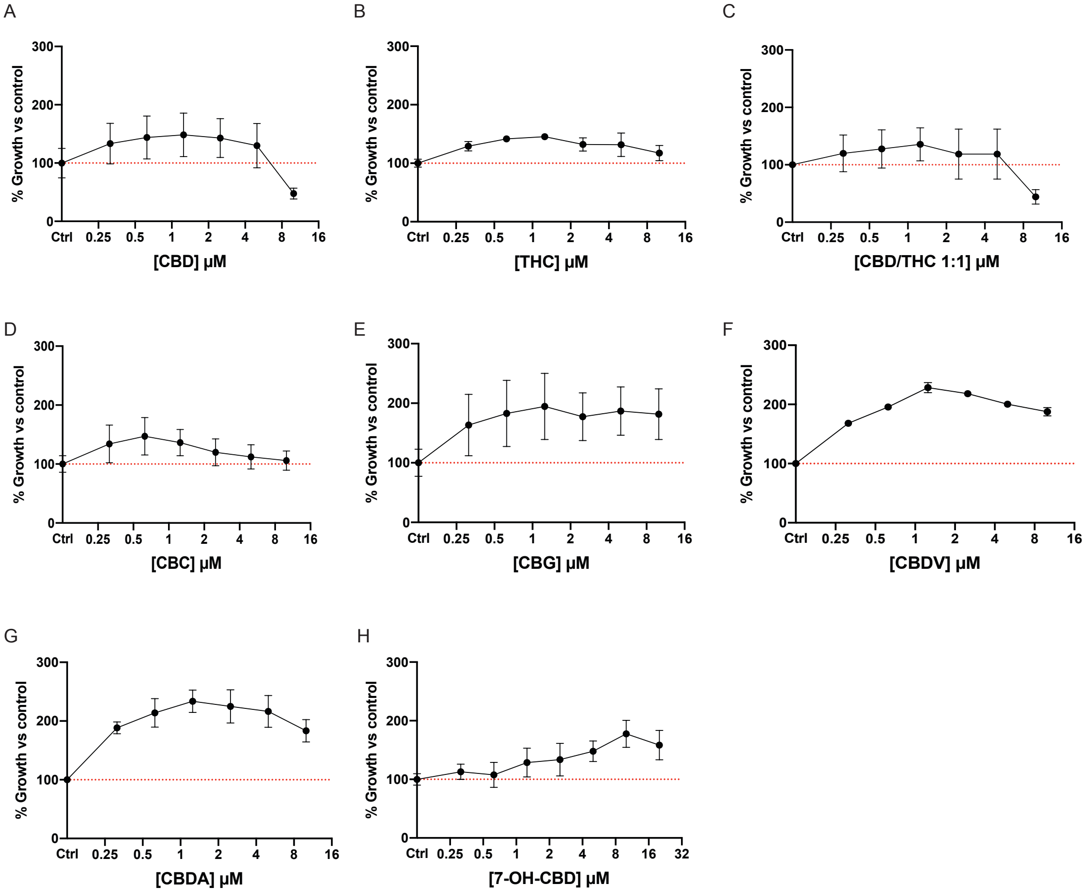


**Fig. S2. Viability of A549-ACE2 cells in different cannabinoids.** A549-ACE2 cells were treated with indicated doses of each cannabinoid for 48 hours. Cell viability was obtained as described in fig. S1B. (A-G) Viability data for cannabinoids used in Fig. 2A are shown. (H) Viability data for 7-OH-CBD used in Fig. 2C is shown.

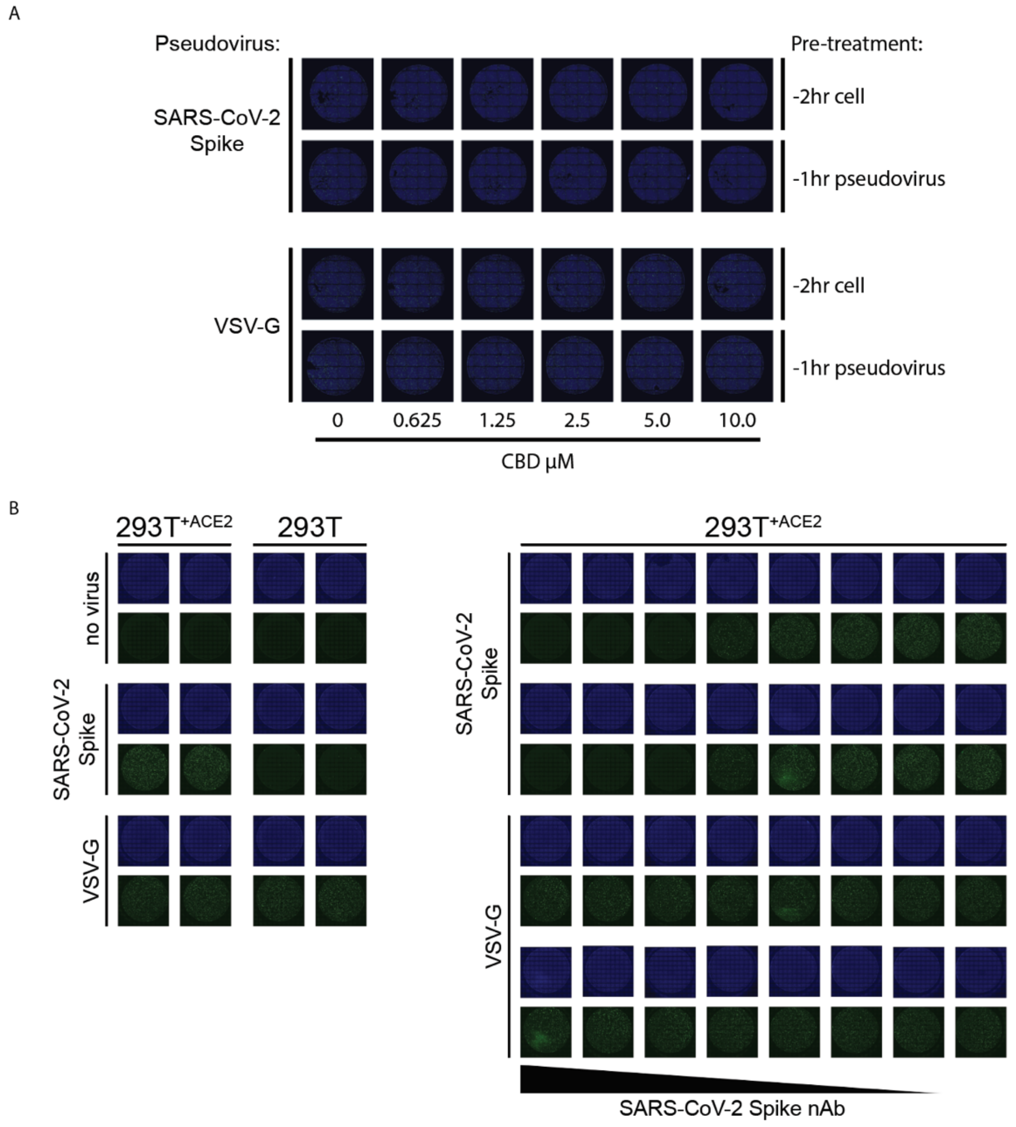


**Fig. S3. CBD's restriction of SARS-CoV-2 infection likely does not occur during entry.** (A) 293T+ACE2 cells were transduced with pseudovirus expressing either SARS-CoV-2 spike or VSV-G at 72hr post-infection following cell or pseudovirus treatment with CBD. Green = Pseudovirus infected, Blue = Nuclear marker Hoechst 33342. (B) 293T or 293T-ACE2 cells were infected for 72hrs with GFP-expressing lentiviruses pseudotyped with either SARS-CoV-2 spike or VSV-G. Pseudoviruses were incubated with 3-fold serial dilutions of SARS-CoV-2 spike neutralizing antibody (highest dose 300ng/100ul/well) for 1hr at 37C prior to infection. Green = Pseudovirus infected, Blue = Nuclear marker Hoechst 33342.

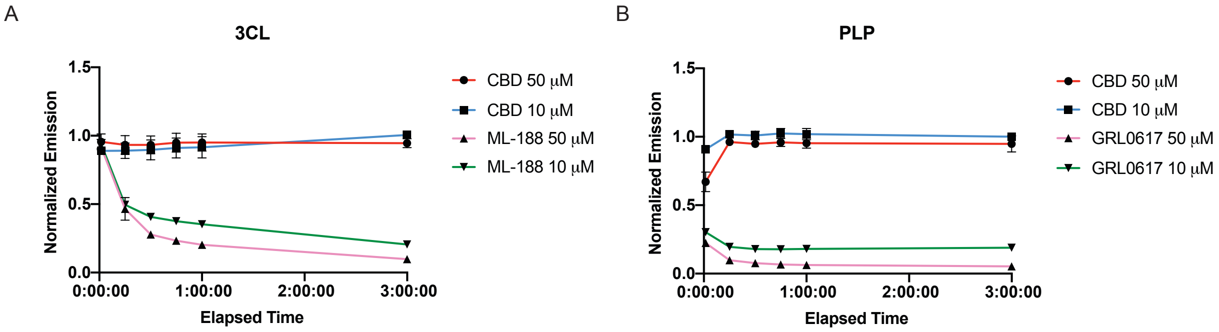


**Fig. S4. CBD does not inhibit the two main viral proteases.** Enzyme assays were performed in duplicate containing indicated concentrations of CBD or positive control inhibitors and 3CLpro enzyme (0.4 μM, A) or PLpro enzyme (0.3 μM, B) as described in Methods. Data at each timepoint were normalized to the negative control (vehicle) wells and the normalized emission levels were plotted.

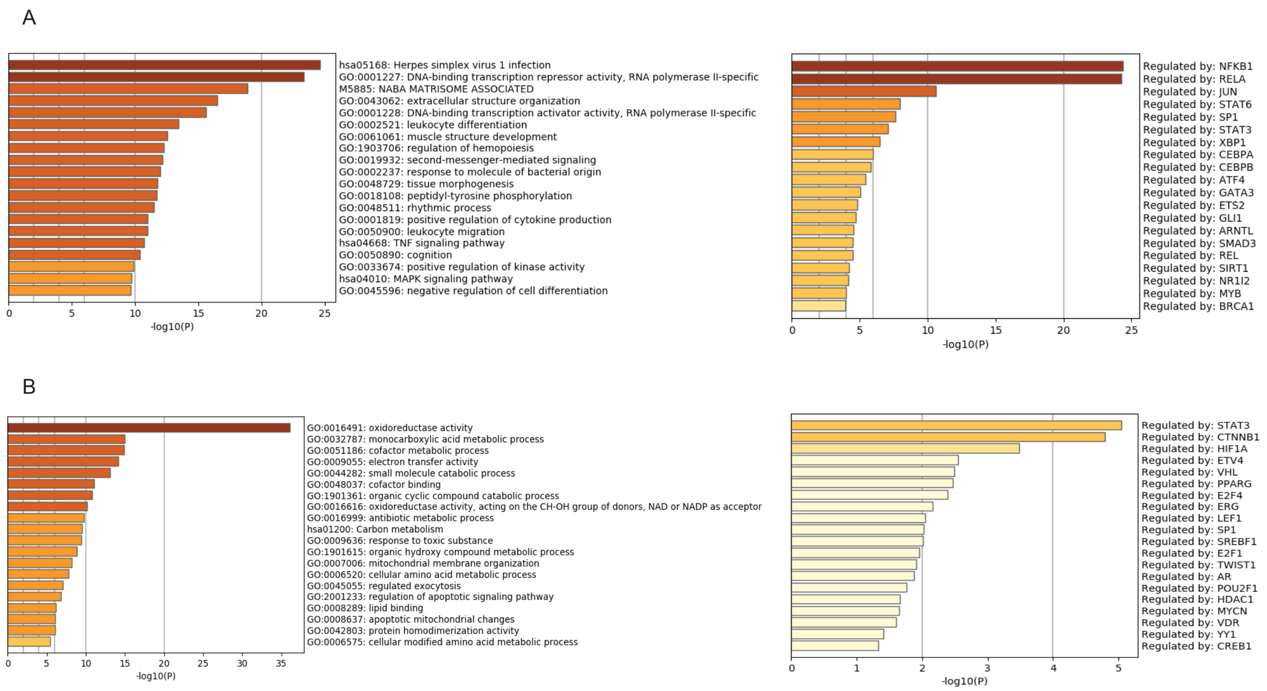


**Fig. S5. Metascape analyses of genes with expression significantly altered by SARS-CoV-2 infection.** Left: Enriched sets of genes with expression significantly up-regulated (A) or down-regulated (B) by SARS-CoV-2 infection from the RNA-seq data were plotted. Differential expression for each gene was determined by the $\log_2$(fold change) of at least 1 in either direction with q-value < 0.01. Right: Possible transcription factors responsible for the transcriptional changes were identified by using the TRUSST database as described in Methods.

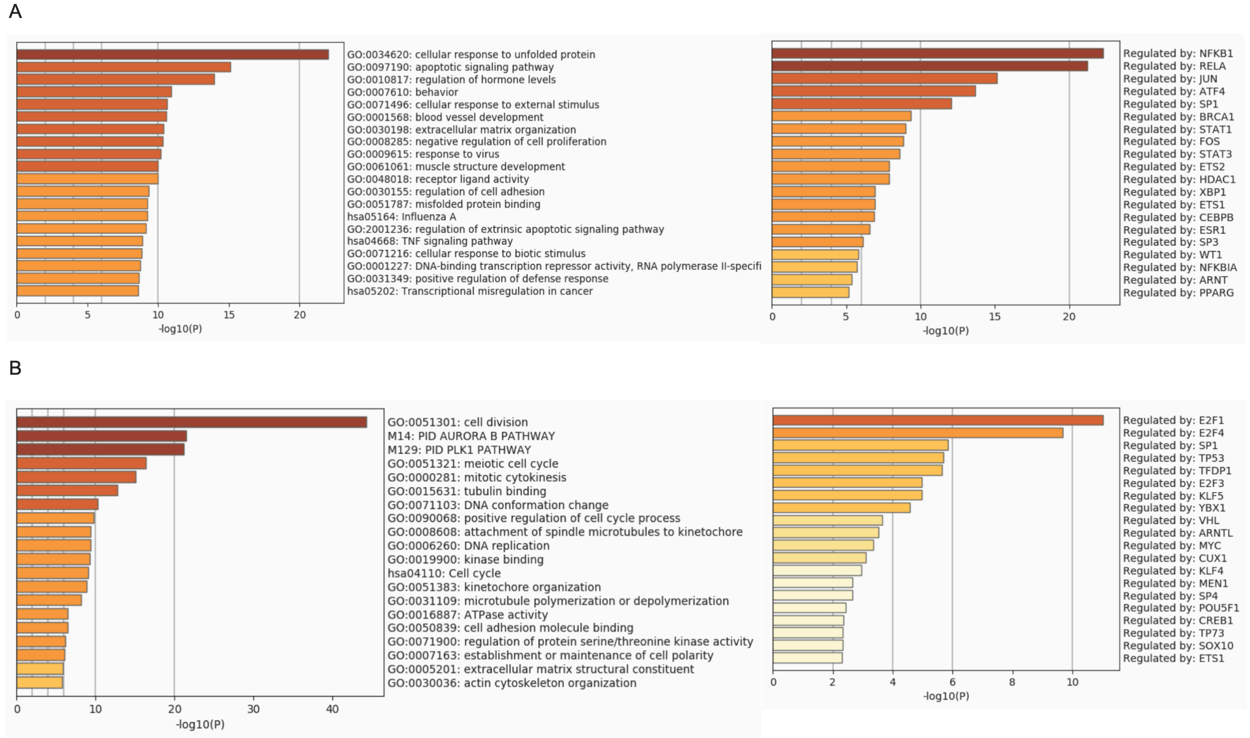


**Fig. S6. Metascape analyses of genes with expression significantly altered by CBD treatment.** Left: Enriched sets of genes with expression significantly up-regulated (A) or down-regulated (B) by CBD treatment from the RNA-seq data were plotted. Differential expression for each gene was determined by the $log_2$(fold change) of at least 1 in either direction with q-value < 0.01. Right: Possible transcription factors responsible for the transcriptional changes were identified by using the TRUSST database as described in Methods.

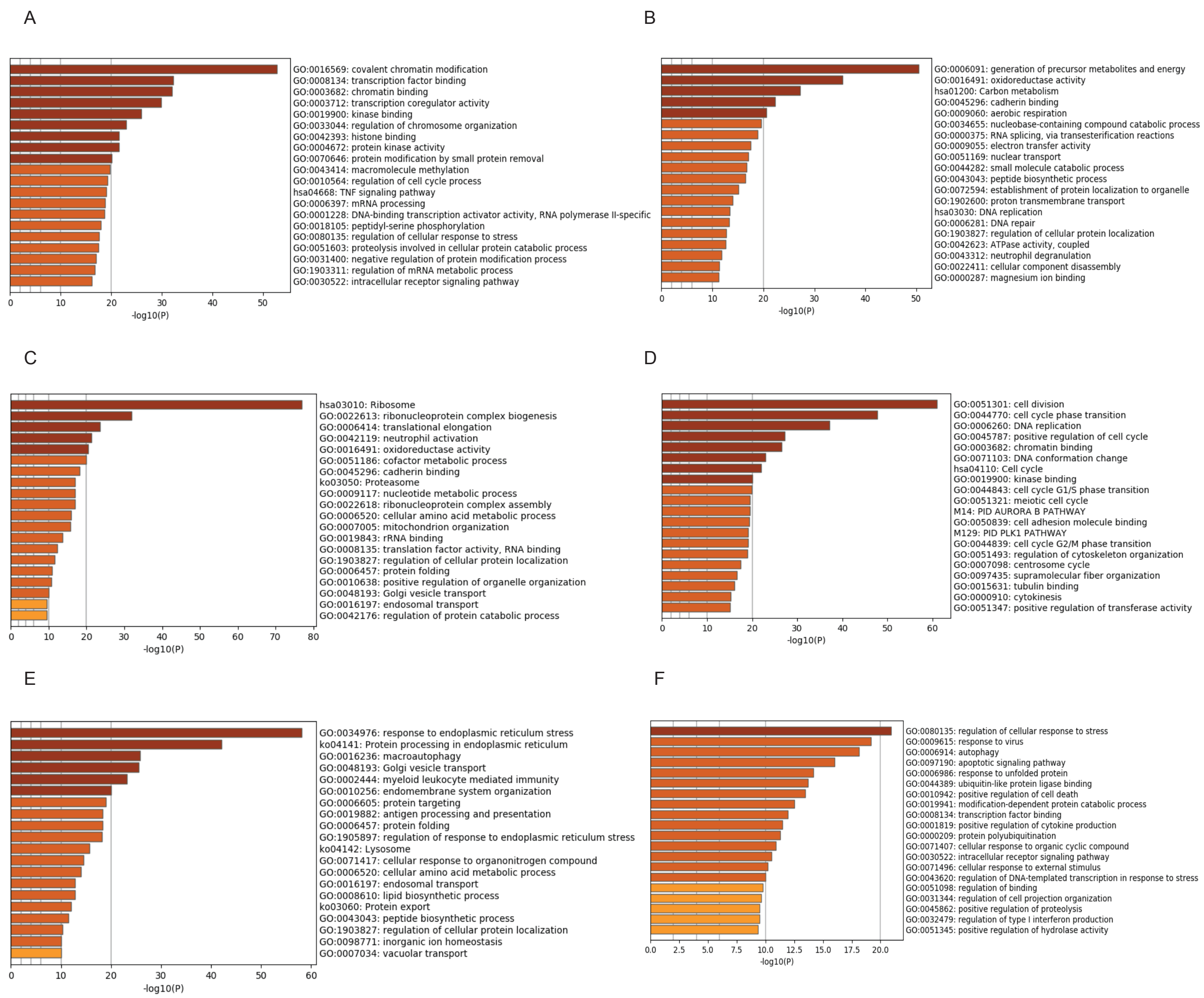


**Fig. S7. Metascape analyses of the most variable genes among the RNA-seq samples.** (A-F) Enriched sets of genes from each of the six clusters as described in Fig. 4D were plotted.

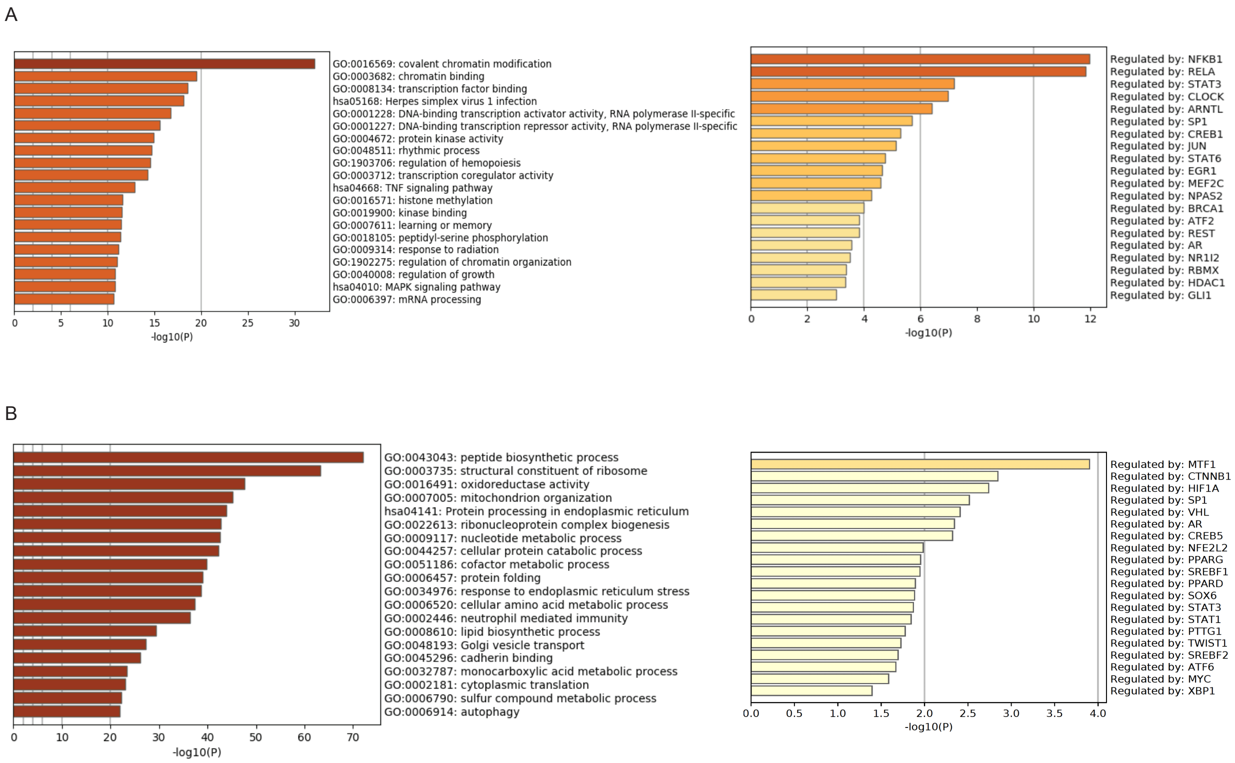


**Fig. S8. Metascape analyses of genes with transcriptional changes due to SARS-CoV-2 infection but reversed by CBD treatment.** Left: Enriched sets of genes with expression significantly up-regulated by viral infection but down-regulated by CBD treatment (A) or vice versa (B) were plotted. Differential expression for each gene was determined by the $\log_2$(fold change) of at least 1 in either direction with q-value < 0.01. Right: Possible transcription factors responsible for the transcriptional changes were identified by using the TRUSST database as described in Methods.

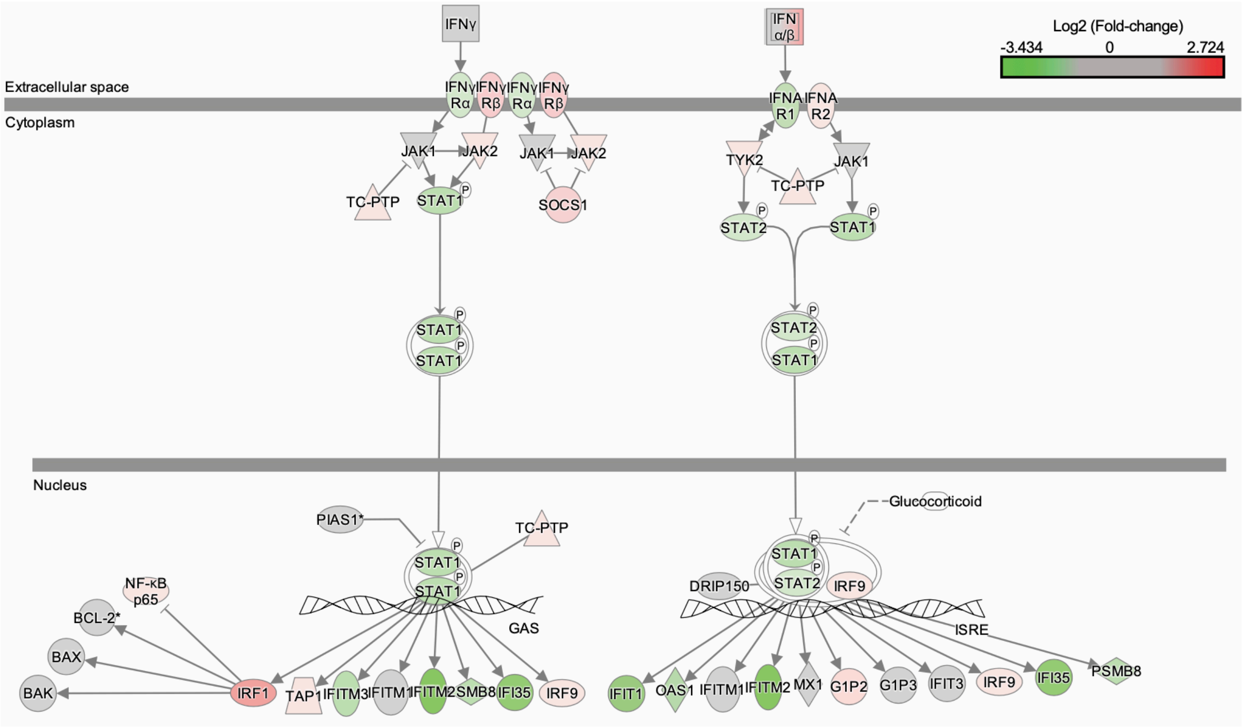


**Fig. S9. Changes in the host interferon signaling canonical pathway following SARS-CoV-2 infection compared to uninfected cells.** In this Ingenuity-generated plot, up-regulated genes are colored red and down-regulated genes are colored green. Color intensity corresponds to $log_2$(fold-change) from RNA-seq.

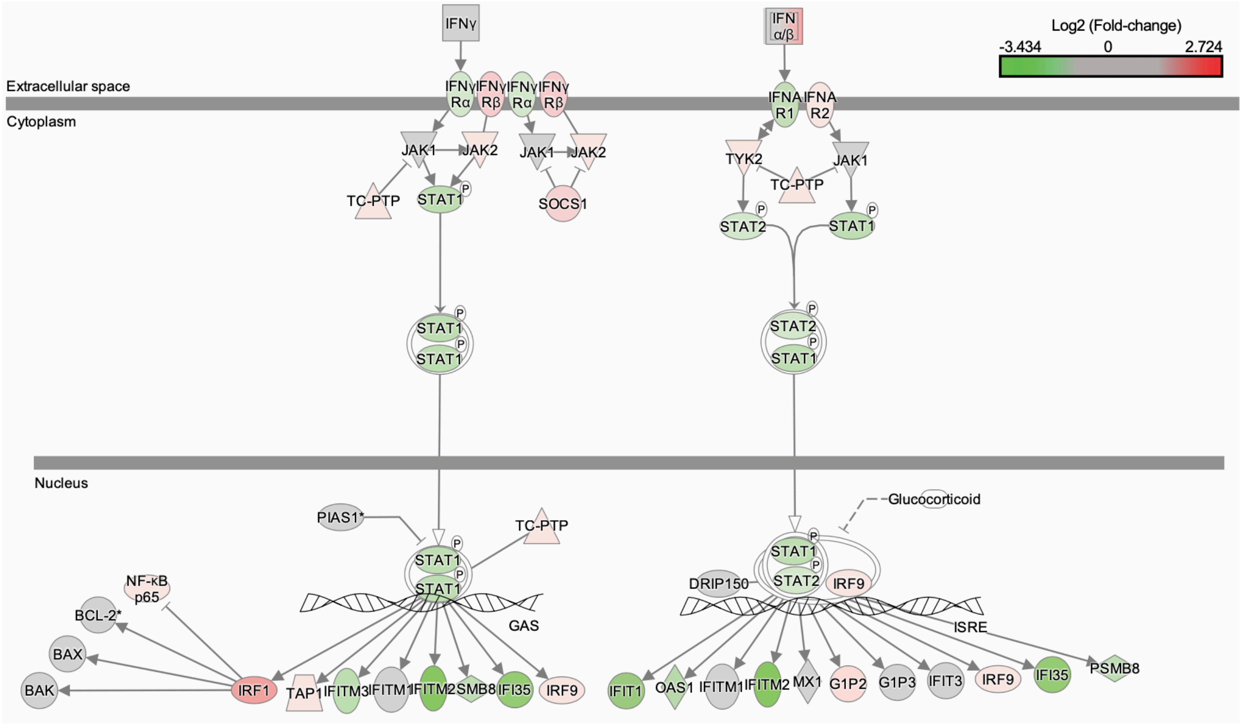


**Fig. S10. Changes in the host interferon signaling canonical pathway with CBD treatment compared to untreated cells.** In this Ingenuity-generated plot, up-regulated genes are colored red and down-regulated genes are colored green. Color intensity corresponds to $log_2$(fold-change) from RNA-seq.

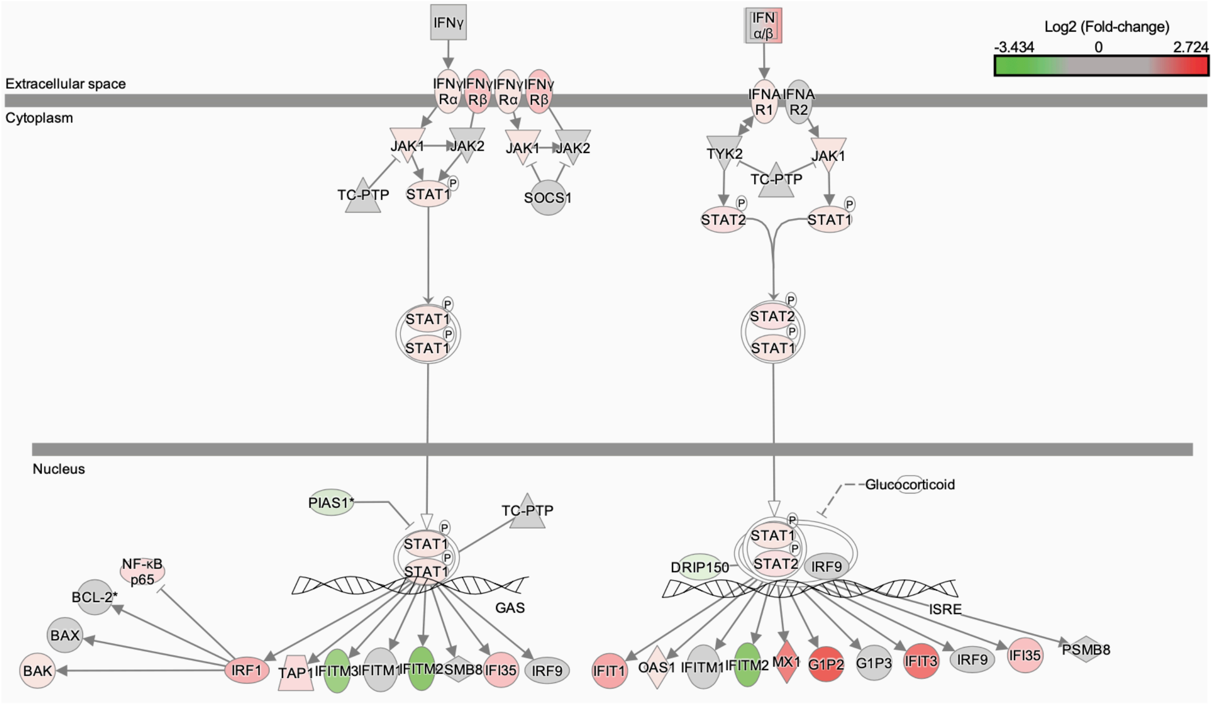


**Fig. S11. Changes in the host interferon signaling canonical pathway following SARS-CoV-2 infection and CBD treatment compared to viral infection alone.** In this Ingenuity-generated plot, up-regulated genes are colored red and down-regulated genes are colored green. Color intensity corresponds to $\log_2$(fold-change) from RNA-seq.

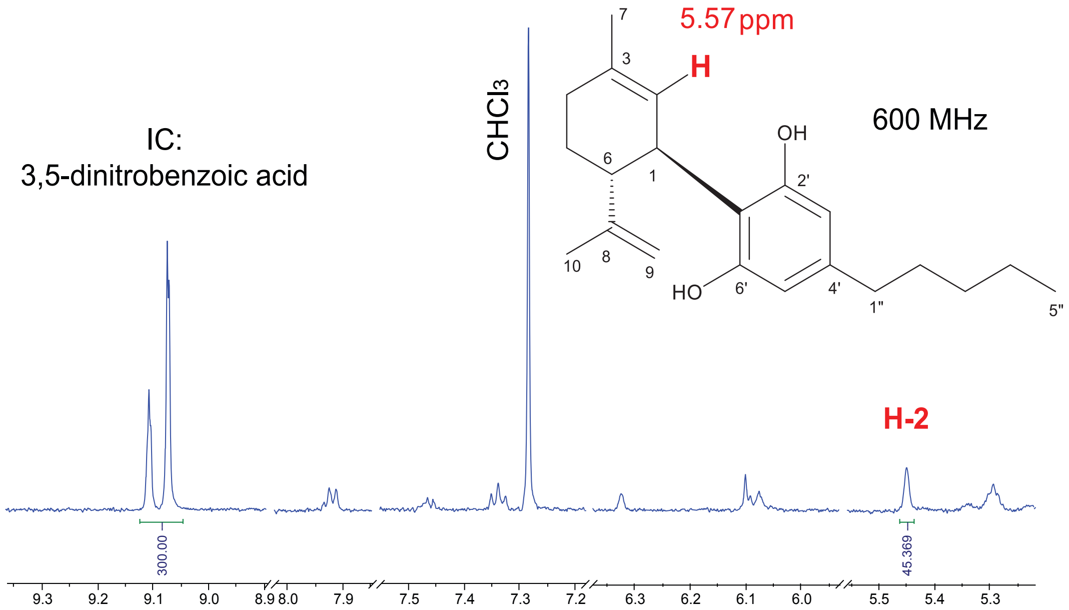


**Fig. S12. $^1$H NMR spectrum of hemp oil.** The absolute qNMR method with internal calibration (3,5-dinitrobenzoic acid as the calibrant) was used to determine the content of CBD in hemp oil as 0.30%. Data was acquired on 600 MHz NMR instrument as described in Methods. Signal at 5.57 ppm (H-2 of CBD) was used for determining CBD quantity.

**Table S1. Characteristics of the patient analysis sample.**

| | Epidiolex Patients[a] (N=82) | | Matched Controls[b] (N=82) | | P-value from $\chi^2$ test |
|---|---|---|---|---|---|
| **Characteristic** | **N** | **%** | **N** | **%** | |
| Age[c] | | | | | |
| 18-29 | 7 | 9 | 5 | 6 | 0.79 |
| 30-39 | 13 | 16 | 15 | 18 | |
| 40-49 | 8 | 10 | 3 | 4 | |
| 50-59 | 14 | 17 | 16 | 20 | |
| 60-69 | 15 | 18 | 16 | 20 | |
| 70-79 | 18 | 22 | 18 | 22 | |
| 80 or greater | 7 | 9 | 9 | 11 | |
| Sex | | | | | |
| Female | 53 | 65 | 48 | 59 | 0.42 |
| Male | 29 | 35 | 34 | 41 | |
| Race | | | | | |
| Black/African-American | 20 | 24 | 26 | 32 | 0.32 |
| White | 51 | 62 | 50 | 61 | |
| Other[d] | 11 | 13 | 6 | 7 | |
| Marital Status[e] | | | | | |
| Married | 40 | 49 | 37 | 45 | 0.64 |
| Not Married | 42 | 51 | 45 | 55 | |
| Religion | | | | | |
| Other[f] | 34 | 41 | 36 | 44 | 0.71 |
| Roman Catholic | 12 | 15 | 13 | 16 | |
| Unknown | 21 | 26 | 15 | 18 | |
| None | 15 | 18 | 18 | 22 | |
| Morbidities | | | | | |
| Conditions with immunosuppression | 41 | 50 | 33 | 40 | 0.21 |
| Hypertension | 37 | 45 | 42 | 51 | 0.44 |
| Chronic lung disease | 21 | 26 | 24 | 29 | 0.60 |
| Depression | 20 | 24 | 19 | 23 | 0.85 |
| Neurologic, seizures | 11 | 13 | 14 | 17 | 0.52 |
| Liver disease | 11 | 13 | 9 | 11 | 0.63 |
| Neurologic, other | 10 | 12 | 12 | 15 | 0.65 |
| Diabetes | 10 | 12 | 8 | 10 | 0.62 |
| Renal failure | 6 | 7 | 10 | 12 | 0.29 |
| Neurologic, movement | 4 | 5 | 4 | 5 | 1.00 |
| Psychoses | 3 | 4 | 3 | 4 | 1.00 |
| Pulmonary circulatory disorders | 3 | 4 | 5 | 6 | 0.47 |
| Dementia | 1 | 1 | 2 | 2 | 0.56 |
| Medication category[g] | | | | | |

| | Epidiolex Patients[a] (N=82) | | Matched Controls[b] (N=82) | | P-value from $\chi^2$ test |
|---|---|---|---|---|---|
| **Characteristic** | **N** | **%** | **N** | **%** | |
| sodium | 53 | 65 | 56 | 68 | 0.62 |
| acetam | 48 | 59 | 52 | 63 | 0.52 |
| lidoca | 46 | 56 | 47 | 57 | 0.88 |
| hydroc | 44 | 54 | 45 | 55 | 0.88 |
| fentan | 44 | 54 | 41 | 50 | 0.64 |
| ondans | 41 | 50 | 39 | 48 | 0.76 |
| midazo | 41 | 50 | 35 | 43 | 0.35 |
| lactat | 38 | 46 | 37 | 45 | 0.88 |
| prochl | 35 | 43 | 30 | 37 | 0.43 |
| propof | 34 | 41 | 33 | 40 | 0.87 |
| hydrom | 31 | 38 | 28 | 34 | 0.63 |
| gabape | 31 | 38 | 38 | 46 | 0.27 |
| hepari | 29 | 35 | 31 | 38 | 0.75 |
| magnes | 29 | 35 | 27 | 33 | 0.74 |
| tramad | 28 | 34 | 33 | 40 | 0.42 |
| ibupro | 28 | 34 | 33 | 40 | 0.42 |
| loraze | 27 | 33 | 29 | 35 | 0.74 |
| trimet | 27 | 33 | 24 | 29 | 0.61 |
| diphen | 26 | 32 | 28 | 34 | 0.74 |
| dexame | 26 | 32 | 25 | 30 | 0.87 |
| potass | 26 | 32 | 24 | 29 | 0.73 |
| pantop | 26 | 32 | 25 | 30 | 0.87 |
| phenyl | 26 | 32 | 23 | 28 | 0.61 |
| flutic | 25 | 30 | 24 | 29 | 0.87 |
| dextro | 24 | 29 | 17 | 21 | 0.21 |
| bupiva | 24 | 29 | 28 | 34 | 0.50 |
| albute | 23 | 28 | 24 | 29 | 0.86 |
| rocuro | 23 | 28 | 24 | 29 | 0.86 |
| sugamm | 23 | 28 | 24 | 29 | 0.86 |
| oxycod | 22 | 27 | 23 | 28 | 0.86 |
| sennos | 21 | 26 | 23 | 28 | 0.72 |
| vancom | 20 | 24 | 19 | 23 | 0.85 |
| enoxap | 20 | 24 | 23 | 28 | 0.59 |
| polyet | 20 | 24 | 21 | 26 | 0.86 |
| cefazo | 19 | 23 | 20 | 24 | 0.85 |
| docusa | 19 | 23 | 18 | 22 | 0.85 |
| generi | 19 | 23 | 24 | 29 | 0.38 |
| calciu | 18 | 22 | 20 | 24 | 0.71 |
| furose | 17 | 21 | 19 | 23 | 0.71 |
| famoti | 16 | 20 | 19 | 23 | 0.57 |
| triamc | 15 | 18 | 13 | 16 | 0.68 |

| | Epidiolex Patients[a] (N=82) | | Matched Controls[b] (N=82) | | P-value from $\chi^2$ test |
|---|---|---|---|---|---|
| **Characteristic** | **N** | **%** | **N** | **%** | |
| naloxo | 15 | 18 | 17 | 21 | 0.69 |
| metocl | 14 | 17 | 10 | 12 | 0.38 |
| ketoro | 14 | 17 | 14 | 17 | 1.00 |
| insuli | 14 | 17 | 13 | 16 | 0.83 |
| predni | 13 | 16 | 15 | 18 | 0.68 |
| entera | 13 | 16 | 15 | 18 | 0.68 |
| epinep | 12 | 15 | 7 | 9 | 0.22 |
| metron | 12 | 15 | 10 | 12 | 0.65 |
| methyl | 12 | 15 | 7 | 9 | 0.22 |
| duloxe | 12 | 15 | 17 | 21 | 0.31 |
| morphi | 11 | 13 | 12 | 15 | 0.82 |
| metopr | 11 | 13 | 11 | 13 | 1.00 |
| cefepi | 8 | 10 | 7 | 9 | 0.79 |
| montel | 7 | 9 | 11 | 13 | 0.32 |
| fat-em | 7 | 9 | 2 | 2 | 0.09 |
| pn-adu | 6 | 7 | 2 | 2 | 0.15 |
| hydrox | 6 | 7 | 6 | 7 | 1.00 |
| pegfil | 5 | 6 | 1 | 1 | 0.10 |
| altepl | 5 | 6 | 4 | 5 | 0.73 |
| fosapr | 5 | 6 | 3 | 4 | 0.47 |
| divalp | 4 | 5 | 7 | 9 | 0.35 |
| cispla | 3 | 4 | 1 | 1 | 0.31 |
| paclit | 3 | 4 | 2 | 2 | 0.65 |
| gemcit | 3 | 4 | 1 | 1 | 0.31 |
| phenyt | 2 | 2 | 3 | 4 | 0.65 |
| etopos | 2 | 2 | 1 | 1 | 0.56 |
| azacit | 1 | 1 | 2 | 2 | 0.56 |
| nab-pa | 1 | 1 | 0 | 0 | 0.32 |
| natali | 1 | 1 | 0 | 0 | 0.32 |

[a] There were 93,565 distinct adults tested for COVID-19 at UCM between March 3, 2020 and January 19, 2021. 85 (<0.1%) had a CBD medication started before their COVID-19 test order date, of whom 82 had started Epidiolex® and were therefore labeled Epidiolex Patients.

[b] Matched Controls were selected algorithmically from Feasible Controls. In detail, Feasible Controls are patients age 18 or older with no prescribed or self-reported cannabinoid use, complete data for all characteristics in this table, and at least medication documented in the University of Chicago Medicine electronic medical record (UCM EMR) in 15 to 730 days before their first COVID-19 test order date. 40,544 (43%) of the 93,565 patients were Feasible Controls. 52,506 (99%) of the 52,936 infeasible patients were considered infeasible because they had no medication documented in the UCM EMR 15 to 730 days before their first COVID-19 test order date and were therefore

potentially more likely to have unobserved cannabinoid use, 116 (0.2%) because they had documentation of a non-Epidiolex® cannabinoid in the UCM EMR 15 to 730 days before their first COVID-19 test order date, and the remaining 314 (0.6%) because of missing responses to age, race, marital status, or religion questions. Matched Controls are Feasible Controls who had the closest propensity score to a patient in the Epidiolex Patients group, where propensity scores were calculated by a logistic regression model fitted to all Epidiolex Patients and feasible controls using the characteristics shown in the table. See the statistical analysis subsection of the Methods and Materials for more detail about the matching algorithm. The pseudo-R2 for the propensity score model was 0.224, the area under its receiver operating characteristic curve was 0.874, and the mean (standard deviation) propensity scores was 0.060 (0.122) for the Epidiolex Patients and 0.053 (0.097) for the Matched Controls (two-sided t-test p-value = 0.70).

[c] The mean (standard deviation) age was 56.5 (18.8) for Epidiolex Patients and 58.2 (18.5) for Matched Controls (two-sided t-test p-value = .56).

[d] The race category Other includes 7 patients reporting “More than one Race,” 5 reporting “Asian/Mideast Indian,” 2 reporting Unknown, 2 reporting Declined, and 1 reporting American Indian or Native American. Otherwise, patients reported “Black/African-American” or “White.” For ethnicity, 149 patients reported “Not Hispanic or Latino,” 12 “Hispanic or Latino,” 2 Declined, and 1 Unknown. 7 of the 12 patients reporting Hispanic or Latino reported a race in the Other race category.

[e] The marital status category Other included 79 patients who reported being single, divorced, legally separated, or widowed, and 8 patients who reported having unknown marital status (4 Epidiolex® Patients, 4 controls). The Married category included 76 patients who reported being married and 1 who reported having a lifetime partner.

[f] 54 of the 70 religion category Other reported non-Roman-Catholic Christian denominations.

[g] All non-cannabinoid medications were categorized by the first six characters of their medication names, which are defined in the UCM EMR.
